# A mathematical model of calcium signals around laser-induced epithelial wounds

**DOI:** 10.1101/2022.08.24.505161

**Authors:** Aaron C. Stevens, James T. O’Connor, Andrew D. Pumford, Andrea Page-McCaw, M. Shane Hutson

## Abstract

Cells around epithelial wounds must first become aware of the wound’s presence in order to initiate the wound healing process. An initial response to an epithelial wound is an increase in cytosolic calcium followed by complex calcium signaling events. While these calcium signals are driven by both physical and chemical wound responses, cells around the wound will all be equipped with the same cellular components to produce and interact with the calcium signals. Here, we have developed a mathematical model in the context of laser-ablation of the *Drosophila* pupal notum that integrates tissue-level damage models with a cellular calcium signaling toolkit. The model replicates experiments in the contexts of control wounds as well as knockdowns of specific cellular components, but it also provides new insights that are not easily accessible experimentally. The model suggests that cell-cell variability is necessary to produce calcium signaling events observed in experiments, it quantifies calcium concentrations during wound-induced signaling events, and it shows that intercellular transfer of the molecule IP_3_ is required to coordinate calcium signals across distal cells around the wound. The mathematical model developed here serves as a framework for quantitative studies in both wound signaling and calcium signaling in the *Drosophila* system.

## 1 Introduction

Wound healing is an essential mechanism that allows an epithelial tissue to maintain its barrier function (Honda, 2017; Martini *et al*., 2017). As a first step, cells must be alerted to the presence of a nearby wound (Stevens and Page-McCaw, 2012; Enyedi and Niethammer, 2015; Park *et al*., 2017). Among early wound-induced signals is an increase in cytosolic calcium – a “life and death signal” that is essential for a wide variety of biological processes (Berridge *et al*., 1998; Xu and Chisholm, 2011; Antunes *et al*., 2013; Razzell *et al*., 2013). Previous work has shown that multiple physical and chemical mechanisms initiate calcium signals around epithelial wounds, some starting as fast as milliseconds, others reaching more distal cells in minutes (Shannon *et al*., 2017; O’Connor *et al*., 2021b). Here, we develop a mathematical model for wound-induced calcium signaling across an epithelium of thousands of coupled cells. The model incorporates two key triggering mechanisms – plasma membrane damage and protease-based activation of latent pro-peptides – which then regulate calcium fluxes across cells’ plasma membranes, between cells’ cytosol and intracellular endoplasmic reticulum (ER) stores, and through gap junctions with neighboring cells. The resulting model yields rich calcium-signaling dynamics that reproduce those observed in experiments, recreate the results of several genetic perturbations, and provide insights not directly accessible through experiments.

The mechanisms included here are those observed using the genetically-encoded calcium indicator GCaMP6m to follow calcium signals around laser wounds in the *Drosophila* pupal notum. Those experiments showed that the initial calcium influx begins within milliseconds and is initiated by cavitation-induced shear stresses that physically create micro-tears in the cells’ plasma membranes. The radius of this damaged region ranged from 30-120 μm, depending on laser energy, and corresponded to 70-1000 cells (not all of which will survive). Over the next 10-20 s, this initial influx spreads through one to two rows of surrounding undamaged cells (O’Connor *et al*., 2021a)– a process blocked by knocking down gap junctions (Shannon *et al*., 2017). In parallel, proteases released by cell lysis near the wound cleave and activate cytokines called growth blocking peptides (Gbps). The activated Gbps diffuse through the extracellular space and bind to the cell surface receptor Methuselah-like 10 (Mthl10), a G-protein coupled receptor, to trigger a canonical G_αq_PLCβ/IPR cascade that releases calcium from the ER. These distal calcium signals appear ~40 s after wounding and spread as far as 150 μm from the wound (O’Connor *et al*., 2021b). Stochastic calcium waves or flares then extend even further away from the wound.

These separate damage events drive different aspects of intra- and intercellular calcium signaling. To understand how they are integrated into a collective tissue-level wound response, we implement the damage signals as inputs into an ordinary differential equation (ODE) model for calcium signaling that is operable in every cell in the epithelium – generating a large set of coupled ODEs. Calcium signaling has been previously modeled in a wide range of systems, from the excitation of muscles and neurons to non-excitable cells such as hepatocytes and *Xenopus* oocytes, and from the kinetics of single calcium channels all the way to signal propagation across entire tissues (Atri *et al*., 1993; Dupont *et al*., 2003, 2007; Graupner and Brunel, 2010; Williams *et al*., 2010; Thurley *et al*., 2011). These wide-ranging models nonetheless have a modular design: specific components of a calcium signaling “toolkit” can be included or excluded based on the system in question (Dupont *et al*., 2016). From this toolkit, we include gap-junction fluxes, regulated cytosol-ER fluxes through the IP_3_ receptor and SERCA pumps, and plasma-membrane fluxes including leak currents, store-operated channels (SOCs), the plasma membrane calcium ATPase (PMCA), and a generalized high-capacity pump. On top of this toolkit, we apply a spatial distribution of micro-tears that add another plasma-membrane flux, and we couple in a previously published reaction-diffusion model for protease-driven activation of Gbps, and their binding to cell-surface receptors.

## 2 Methods

Our mathematical model represents the monolayer epithelium of the *Drosophila* pupal notum as a Voronoi mesh of tightly packed cells. Upon wounding, each cell is subjected to a different degree of damage and experiences different time-dependent levels of active Gbp signaling through the G-protein coupled receptor Mthl10. The cellspecific damage and Gbp-Mthl10 signaling then couples into each cell’s common calcium signaling machinery: fluxes across the plasma membrane; fluxes through gap junctions with neighboring cells; and IP_3_-mediated fluxes between the cytoplasm and intracellular ER stores. This coupling of wound-induced damage, extracellular signals, and intracellular signaling pathways gives rise to the rich calcium signaling dynamics observed in experiments. A schematic of the model is shown in Figure 1, and each part is described in detail below.

**Figure 1:**
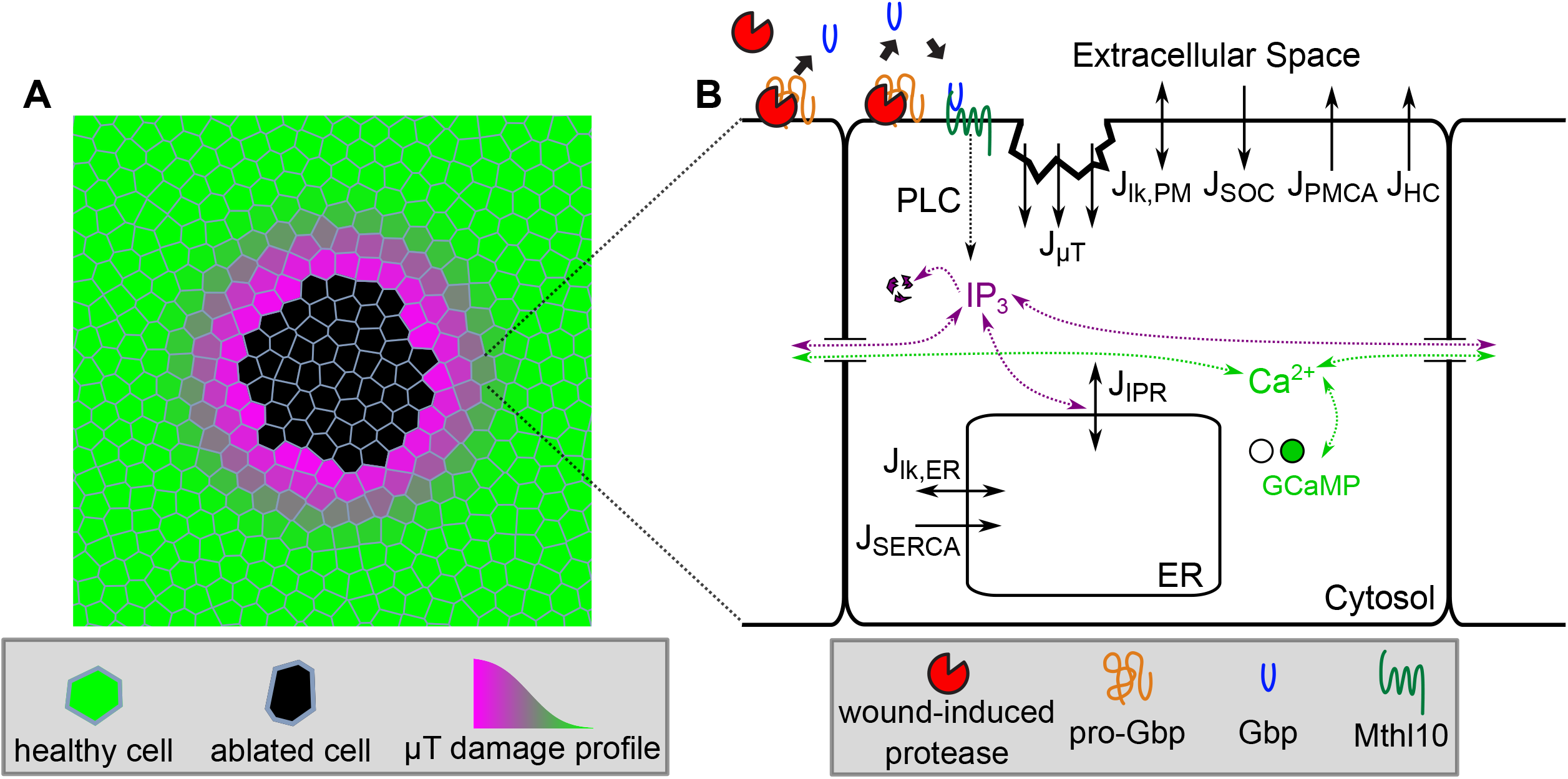
Schematic diagram of the complete wound-induced calcium signaling model. 160 x 160 μm section of the tissue mesh centered around the wound. Cells are color-coded according to their physical damage level. (B) Reactions and fluxes included in the single-cell model and its connection to the extracellular model of biochemical damage signals. Solid black arrows show directions of calcium fluxes.

### 2.1 Tissue-level model

The *in silico* tissue is a 2D Voronoi mesh constructed from a set of 4826 seed points confined to a 450 μm by 450 μm square. The seed points were initially placed at random and then redistributed more uniformly by simulated annealing with an inverse-square-distance repulsion between all seed pairs. A similar force repulsed each seed from the closest boundary of the square. After Voronoi tessellation, each Voronoi cell represents one modeled biological cell. Compared to polygonal approximations of segmented experimental images, these Voronoi meshes have similar means for cell area, cell perimeter, and cell packing (number of neighbors); however, the distributions for the model meshes are narrower and have smaller standard deviations (Figure S1). A 160 μm by 160 μm section of an example tissue mesh is shown in Figure 1A.

### 2.2 Wound-induced damage signals

#### 2.2.1 Plasma Membrane Micro-tears

The wounds we model are those caused by pulsed laser ablation (Shannon *et al*., 2017). Such wounds locally destroy some cells – those labeled as “ablated” in Figure 1A – and permeabilize others further out through damage to their plasma membranes, so called micro-tears. This damage is caused by shear stress created through the rapid (< 100 μs) expansion and collapse of a laser-induced cavitation bubble (Vogel and Venugopalan, 2003; Hellman *et al*., 2007). We previously found that the radius of the region of calcium influx at ~2 s after wounding was well-matched to the maximum bubble radius. By imaging even faster, we find that a central region of calcium influx still appears within the first frame (17 ms after wounding), but more distal influx proceeds from discrete sites that fill in the cavitation-bubble footprint over 1-2 seconds.

Analysis of the density of discrete calcium entry sites provides an estimate for the distribution of plasma-membrane damage (Figure S2). We find that this distribution can be heuristically described by

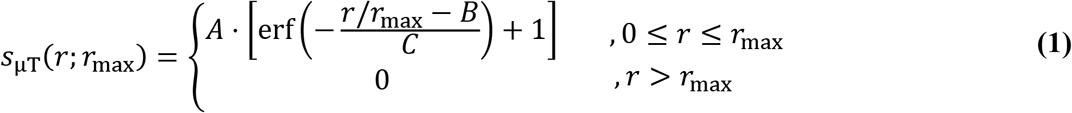

where *s*_μT_ is the local plasma-membrane permeability due to micro-tears, erf is the error function 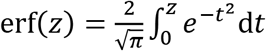 is the maximum radius of the cavitation bubble, and *A, B*, and *C* are fitting parameters with values given in Table 1. This heuristic function provides a means to scale the damage distribution for wounds with different sized cavitation bubbles.

**Table 1:** Parameter values for the damage models.

| Symbol | Definition | Value |
| --- | --- | --- |
| *Plasma Membrane Damage Distribution* |  |  |
| $A$ | Damage distribution parameter | 0.48 |
| $B$ | Damage distribution parameter | 0.72 |
| $C$ | Damage distribution parameter | 0.16 |
| $r_{\max}$ | Maximum cavitation bubble radius | 51 $\mu\text{m}$ |
| *Extracellular GBP-Mthl10 Signaling* |  |  |
| $\gamma_{DP}$ | Scaled protease diffusion constant | 0.012 |
| $\gamma_{DL}$ | Scaled Gbp diffusion constant | 134 |
| $\gamma_s$ | Scaled protease release rate | 0.019 |
| $\gamma_c$ | Scaled protease cleavage rate | 487 |
| $\gamma_P$ | Scaled total protease amount | 1.15 |
| $\gamma_{pL}$ | Scaled total Gbp amount | 0.025 |
| $\gamma_L$ | Scaled total Gbp amount | 1.17 |
| $\gamma_R$ | Scaled total Mthl10 amount | 0.023 |
| $\gamma_w$ | Scaled protease release radius | 2.41 |

The parameters for the plasma membrane damage distribution are determined through fits to data as explained in the text. The parameters for the extracellular GBP-Mthl10 signaling model are described in more detail in O'Connor et al. (2021b), but are obtained assuming a calcium signaling threshold of $w_\ell = 0.05$ rather than 0.5.

#### 2.2.2 Extracellular Gbp-Mthl10 Signaling

After the initial calcium influx, there is a second delayed calcium response driven by an extracellular signaling cascade: proteases released from lysed and damaged cells cleave latent, extracellular proGbps into their active forms, which diffuse in the extracellular space and bind to the cell-surface receptor Mthl10, which initiates calcium release from the ER. This Mthl10 response extends even more distally than the first calcium response caused by damage to the plasma membranes. As done previously, we model that process as a set of coupled reaction-diffusion equations. The scaled equations (Eqs. (2) – (5) below) are solved over time and distance from the wound to determine *w_ℓ_*, the local fraction of Mthl10 receptors that have bound Gbp. A continuous extracellular space is assumed, so there are no boundary conditions imposed at each cell boundary. The independent variables *ρ* and *τ* are scaled distance and time; the dependent variables *x,y_T_*, and *ℓ* represent local, scaled concentrations of protease, proGbp, and active Gbp respectively; the various *γ* parameters are defined in Table 1. At t=0, there is no protease and all Gbp is in the pro-form. A no-flux boundary is imposed at the origin, and all species are clamped to zero sufficiently far from the wound center (r = 1000 μm). Parameters for downstream intracellular signaling (see Table 1) were selected so that a distal calcium response occurs at a threshold of *w_ℓ_* = 0.05. Further details of the parameters and scaled model equations can be found in O’Connor et al. (2021b)

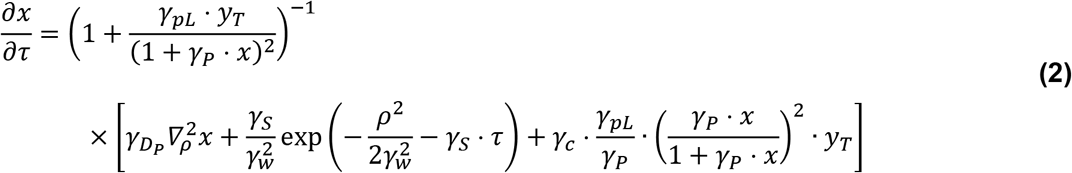

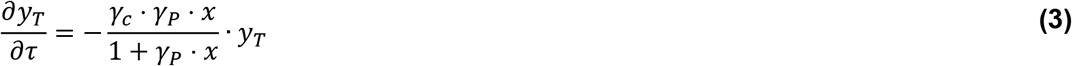

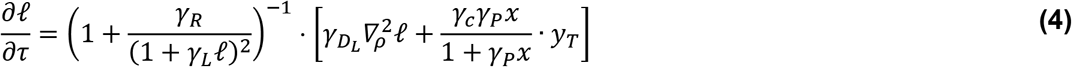

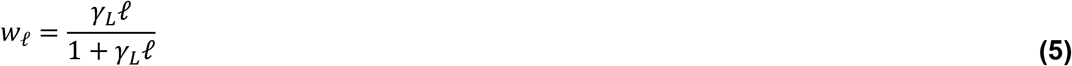

### 2.3 Single-cell model

For each cell in the tissue mesh, the local micro-tear permeability and fraction of Gbp-bound Mthl10 receptors is determined as above and coupled into a single-cell model that uses ordinary, nonlinear differential equations to represent a common set of calcium signaling machinery. The single-cell model tracks cytosolic calcium *c*, ER calcium *c*_ER_, cellular IP_3_ levels *p*, and the IP_3_ Receptor (IPR) inactivation parameter *h*. At the most general level, the ODEs for cytosolic and ER calcium concentrations are based on fluxes and buffering:

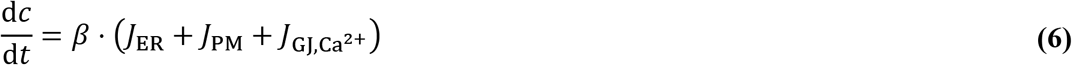

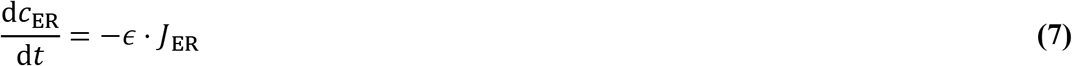

where *J*_ER_, *J*_PM_, and *J*_GJ,Ca^2+^_ are respectively calcium fluxes between a single cell’s cytosol and ER, between a cell’s cytosol and the extracellular space across its plasma membrane, and between adjacent cells through gap junctions. The parameter *ϵ* is the ratio of cytosolic to ER volume, which is needed to express the rate of change of ER calcium concentration in terms of cytosolic calcium fluxes. The variable *β* represents rapid buffering of cytosolic calcium:

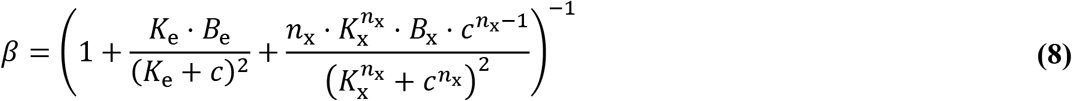

where *K*_e_ and *B*_e_ are the dissociation constant and total concentration of endogenous buffers, and *K*_x_ and *B*_x_ are corresponding parameters for the exogenous, genetically encoded calcium indicator GCaMP6m. Eq. (8) follows from the derivation in Wagner and Keizer (1994) but with the exogenous buffer term modified to reflect GCaMP6m’s experimentally determined Hill coefficient of *n*_x_ = 2.96 (Chen *et al*., 2013).

The value of 5 μM for the total GCaMP concentration (*B*_x_) was chosen in order to be a substantial concentration value without disallowing calcium oscillations completely. Values for the other parameters in this section are taken from various literature sources, and all parameter values and justifications for the single-cell model can be found in Table 2.

**Table 2:**
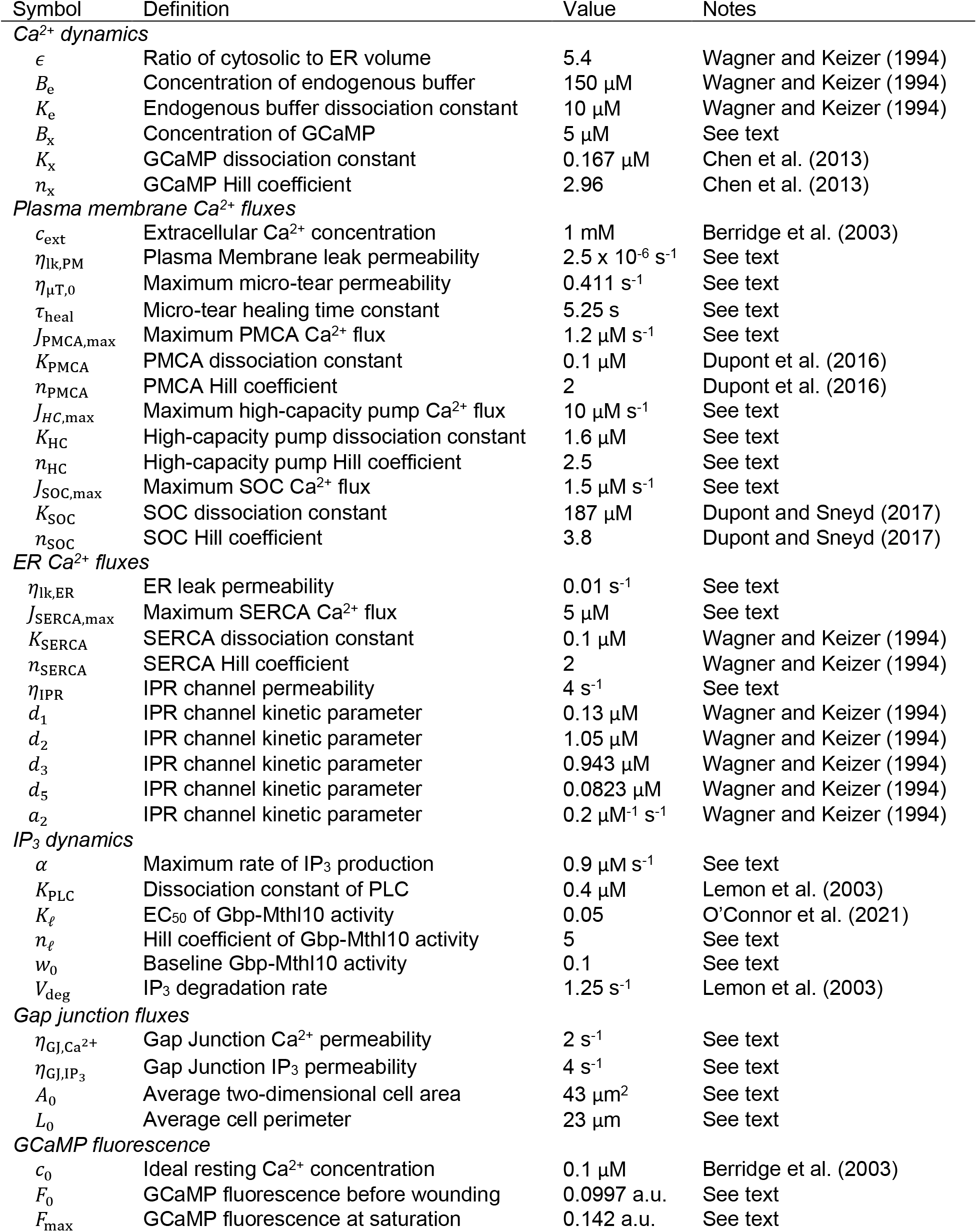
Single-cell model parameter values.

#### 2.3.1 Plasma membrane calcium fluxes

Calcium movement across the plasma membrane of a single cell is modeled by five fluxes: a micro-tear flux present only in damaged cells, a leak flux, and fluxes through store operated channels, PMCA pumps, and a generalized high-capacity pump.

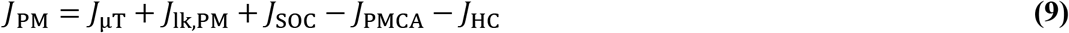

The micro-tear and leak fluxes are modeled as proportional to the difference in calcium concentration across the plasma membrane:

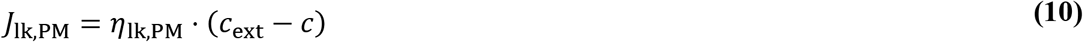

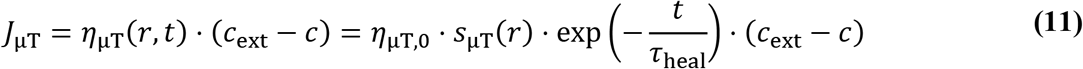

where the leak permeability *η*_lk,PM_ is constant and the initial micro-tear permeability *s*_μT_(*r*) varies with distance from the wound to represent the spatial distribution of membrane damage (see section 2.2.1 and Eq. (1)). The micro-tear permeability decays exponentially with time constant *τ*_heal_ to represent membrane damage repair. For each cell in the tissue mesh, r is the distance from the wound center to the centroid of the cell.

The PMCA and high-capacity pump fluxes are modeled as Hill functions dependent on cytosolic calcium:

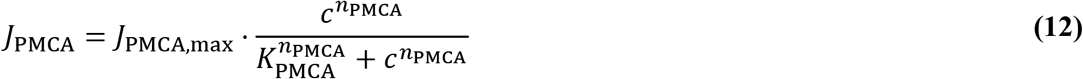

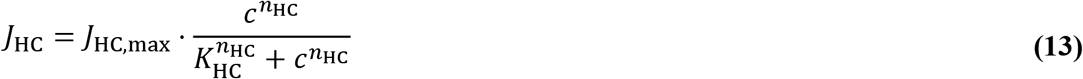

The PMCA is needed to regulate resting calcium levels as well as the calcium flares at lower concentrations, and the high-capacity pump term is needed in order for the first expansion to retract. The model is agnostic as to what the high-capacity pump represents; however, the typical counterpart to the PMCA that is responsible for maintaining cytosolic calcium levels is the sodium-calcium exchanger (NCX) which has been shown to indirectly regulate cell migration in epithelial tissues (Balasubramaniam *et al*., 2015). A complete model of the NCX would require the inclusion of sodium dynamics that are beyond the scope of the model, and so a general calcium-dependent Hill function is used.

The SOC flux is also represented as a Hill function dependent on ER calcium (Dupont and Sneyd, 2017)

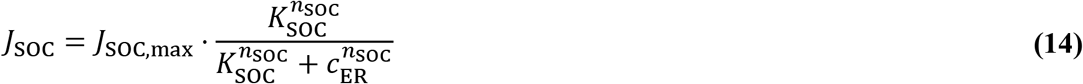

In Eqs. (12) – (14), each *J*_max_ represents a maximum flux value, each *K* represents the calcium concentration at half-maximal activity, and each *n* controls the steepness of the switch from low to high activity. Note that the directionality of these three fluxes is represented directly by the sign preceding each in Eq. (9) (negative for outward fluxes).

Values for parameters *η*_lk,pm_, *J*_PMCA,max_, and *J*_SOC,max_ were chosen to give desired resting cytosolic calcium concentrations (~ 0.1 μM) for single cells as well as to produce propagative calcium waves in the tissue model. Values for parameters *τ*_heal_ and *η*_μT,0_ were chosen such that an isolated cell with a micro-tear permeability of 0.95 · *η*_μT,0_ would reach half of maximum fluorescence in 25 seconds given all other model parameters, consistent with experimental data (not shown). Values for parameters *J*_HC,max_, *K*_HC_, and *n*_HC_ were chosen such that the time and spatial extent of the first expansion matched experimental data.

#### 2.3.2 ER calcium fluxes

Calcium exchange between the cytosol and the ER is determined by three fluxes: a leak flux plus fluxes through the SERCA pump and IP_3_ receptors:

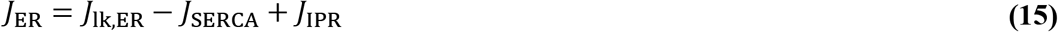

Similar to above, the leak flux simply follows the difference in calcium concentrations between ER and cytosol,

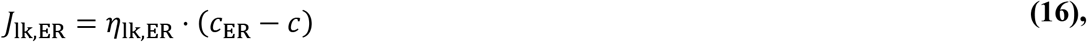

and the SERCA pump is modeled as a Hill function dependent on cytosolic calcium levels,

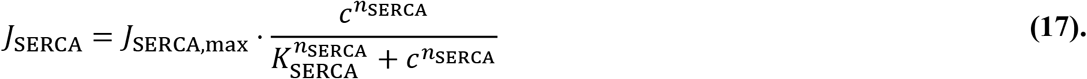

For the IPR flux, we require a deterministic model that involves activation from IP_3_ as well as both activation and inhibition from calcium to produce calcium oscillations. Therefore, we use the relatively simple, effective, and widely used model from Li and Rinzel (1994) for which channel inactivation through low-affinity calcium binding sites is assumed to be much slower than activation through IP_3_ and high-affinity calcium binding. The overall form is

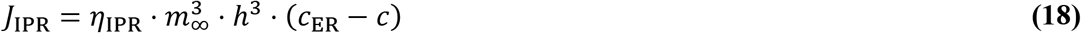

where *m*_∞_ and *h* are respectively fast activation and slow inactivation variables. They are cubed because channel opening requires the coordination of three active IPR subunits. The “fast” activation variable *m*_∞_ depends on cytosolic IP_3_ levels *p* and calcium levels *c*

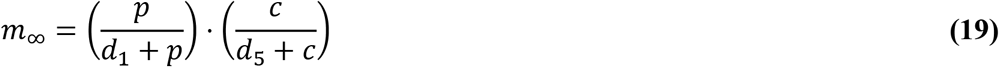

where *d*_1_ and *d*_5_ are the concentrations at which *p* and *c* respectively have half-maximal effects. The “slow” inactivation variable h evolves with time according to

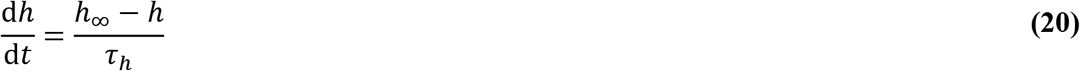

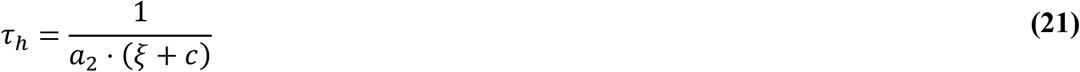

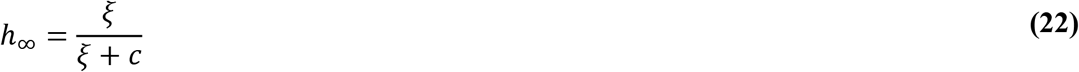

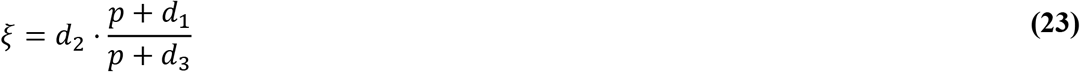

Although more sophisticated models have been published for the IPR flux (Siekmann *et al*., 2012; Cao *et al*., 2014), the Li and Rinzel (1994) model is able to produce calcium oscillations similar to those observed in experiments and is thus sufficient for our system.

Values for parameters *η*_lk,ER_, *J*_SERCA,max_, and *η*_IPR_ were chosen so that resting cytosolic calcium concentrations were ~ 0.1 μM for single cells as well as to produce propagative calcium waves in the tissue model.

#### 2.3.3 IP_3_ dynamics

IP_3_ is generated downstream of Gbp binding to Mthl10, a G-protein-coupled receptor. This is a multi-step process in which ligand-receptor binding activates the G_αq_-protein, which then binds to and activates PLCβ in a calcium-dependent manner, and this complex then cleaves PIP2 into diacylglycerol and IP_3_. Although detailed models for this type of signaling cascade have been published (Lemon *et al*., 2003), we do not have sufficient data to determine those model parameters for our experimental system and have therefore opted for a simpler model of the IP_3_ production rate (*V*_PLC_) that uses a product of Hill functions to represent the cascade’s activation by cytosolic calcium *c* and the fraction of Gbp-bound Mthl10, *w_ℓ_* (as described in section 2.2.2)

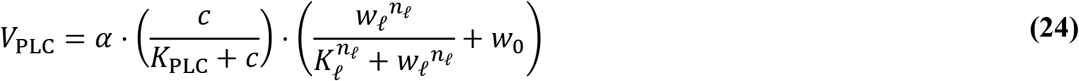

where *α* is the maximum IP_3_ production rate, *n_ℓ_* is a Hill coefficient that controls how sensitive IP_3_ production is to damage-induced Gbp-signaling, and *w*_0_ serves to keep a base-line rate of IP_3_ production in the absence of a wound. Following Lemon et al. (2003), the Hill exponent for the calcium dependence was set to one.

IP_3_ levels also change due to degradation, assumed to be first order, and flux through gap junctions, yielding

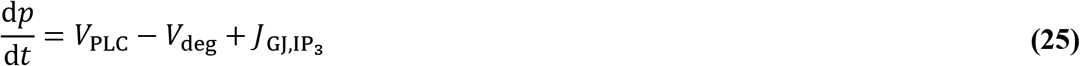

where *p* is the cytosolic concentration of IP_3_, and

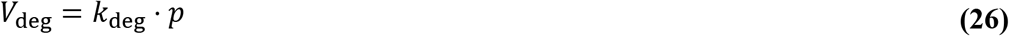

with *k_deg_* constant.

Values for parameters *α, n_ℓ_*, and *w*_0_ were chosen to allow for propagative calcium waves in the tissue-level model as well as to match the time and spatial extent of the distal calcium response to experimental data.

#### 2.3.4 Gap junction transfer of calcium and IP_3_

Adjacent cells are connected by gap junctions that permit intercellular diffusion of both calcium and IP_3_. We use a simple model that does not include any gating effects. Therefore, for cell *j*, the gap junction flux terms for calcium and IP_3_ are given respectively by

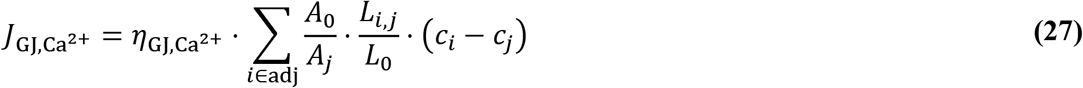

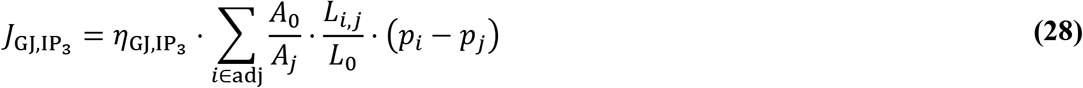

where each sum is over all cells adjacent to cell *j. A_j_* is the 2-dimensional area in the plane of the tissue covered by cell *j*, and *L_i,j_* is the length of the shared edge between cell *i*. and cell *j*. Parameters *A*_0_ and *L*_0_ are a reference area and length respectively, and the ratios *A*_0_/*A_j_* and *L_ij_*/*L*_0_ are explicitly included so that the gap junction permeabilities *η*_GJ,Ca^2+^_ and *η*_GJ,IP_3__ are the same for every cell in the tissue and have comparable dimensions to other *η* parameters in the model corresponding to fluxes due to concentration differences. For cells on the wound margin, a residual gap junction connection to the wound (0.025 · *η*_GJ_) was maintained to reproduce the long-term, high calcium observed around the wound.

The value for parameter *η*_GJ,Ca^2+^_ was chosen to match the time and spatial extent of the first expansion to experimental data. The value for parameter *η*_GJ,IP_3__ was chosen to allow for propagative calcium waves in the tissue model. The value for parameters *A*_0_ and *L*_0_ were measured from segmented tissue images as the average cell area and perimeter respectively; however, the exact values of these parameters are irrelevant; the full gap junction parameter is (*A*_0_/*L*_0_) · *η*_GJ_ for both calcium and IP_3_.

### 2.4 Resting levels and initial conditions

Initial conditions for each cell were determined by running the tissue model with no micro-tears (*η_μT_*(*r*) = 0) and no Mthl10 activation (*w_ℓ_*(*r, t*) = 0) for up to 300 s. Model parameters were selected to ensure that resting cytosolic calcium and IP_3_ concentrations did not deviate too far from 0.1 μM and 10 nM respectively, and that resting ER calcium concentrations were on the order of 100 μM. The cytosolic calcium concentration within the wound was set to be constant and equal to extracellular calcium concentration *c*_ext_ = 1 mM.

### 2.5 GCaMP fluorescence

GCaMP fluorescence *F* is obtained from the model using a Hill function of the free calcium concentration as specified by Chen et al. (2013)

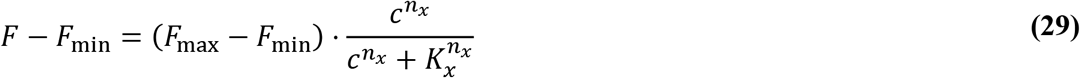

where *F*_min_ is the fluorescence when *c* = 0, and *F*_max_ is the fluorescence at saturation when *c* >> *K_x_*. However, with our *in vivo* system we are not able to determine the GCaMP fluorescence when no calcium is present. Therefore, with the assumption of the resting calcium concentration being *c*_0_ = 0.1 μM, we can express the fluorescence *F* in terms of the resting fluorescence *F*_0_ as

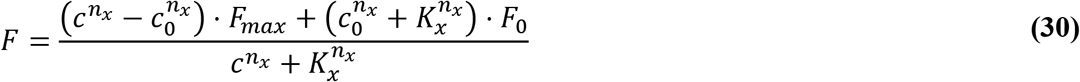

Since GCaMP is a cytosolic reporter, the wound is treated as having no GCaMP, and thus appears black in figures below.

Values for parameters *F*_0_ and *F*_max_ were obtained from GCaMP fluorescence images after wounding from average fluorescence values in regions of resting cytosolic calcium and high cytosolic calcium respectively.

### 2.6 Method of solution

The Gbp-Mthl10 signaling model equations (Eqs. (2) – (5)) are solved numerically in Mathematica over time and over a radially symmetric, continuous space. Since the system of equations is nonlinear and since the equation for proGbp (y_T_) does not contain derivatives with respect to the scaled distance from the wound (ρ), solutions were found using the method of lines. The effectively one-dimensional spatial region was discretized using the “Tensor Product Grid” method. The number of discretized grid points was chosen as the smallest value for which a finer discretization did not yield a significantly different solution. Further details on the method of solution can be found in O’Connor et al. (2021b).

The single-cell model solutions were also solved using Mathematica software. Because these equations are ordinary differential equations, Mathematica employs an LSODA approach, switching between a non-stiff Adams method and a stiff Gear backward differentiation formula method. The initial conditions for each cell in the single-cell model are obtained by running the model for an extended time with no wound (*η*_μT,0_ = 0 and *w_ℓ_* = 0).

All numerical solutions of the model equations are performed in Mathematica on the ACCRE cluster at Vanderbilt University. A single model run along with output video production takes roughly 1.5 to 2.5 hours.

Code and relevant files have been deposited on GitHub (https://github.com/mshutson/wound-calcium-LRCa).

### 2.7 Experimental Methods

Model results are compared to experiments in which laser wounds were made to the notum of *Drosophila* pupae. We summarize the experimental methods below; full details can be found in O’Connor et al (2021 b).

Briefly, white prepupae were identified and aged for 12-18 hours After Puparium Formation (APF) at 29 °C. Each pupa had their anterior pupal case removed with fine tipped forceps to reveal the head and thorax and then adhered to a 35 mm x 50 mm coverslip (Fisherbrand, cat#125485R), notum-side down on a piece of double-sided tape (Scotch brand, catalog #665), thus sandwiching the pupa between the coverslip and tape layer. Then, an oxygen permeable membrane (YSI, standard membrane kit, cat#1329882) was applied over the pupa and secured to the coverslip with additional double-sided tape so the pupa would not become dehydrated or deprived of oxygen.

Laser ablation and live imaging of mounted pupae were performed using a Zeiss LSM410 raster-scanning inverted confocal microscope with a 40X 1.3 NA oil-immersion objective. Laser wounding used single pulses of the third harmonic (355 nm) from a Q-switched Nd:YAG laser (5 ns pulse width; Continuum Minilite II, Santa Clara, CA) at pulse energies on the order of 1 μJ. A separate computer-controlled mirror and custom ImageJ plug-in were used to aim and operate the ablation laser so that ablation could be performed without any interruption to live imaging.

## 3 Results

### 3.1 Model Validation

#### 3.1.1 Control calcium signaling dynamics

Applying Eqs. (1) – (5) across the entire tissue and Eqs. (6) – (28) for each cell in the tissue, the model can calculate cytosolic calcium concentration as a function of time for each cell in the tissue. Applying Eq. (30) then yields expected GCaMP fluorescence for direct comparison to experiments. Using the parameters in Table 2 for every cell, the model yields uniform resting levels of cytosolic calcium, ER calcium, and IP_3_ of 0.09 μm, 220 μM and 13 nm, respectively. These match or are on the same order of magnitude as expected ranges: 0.09 ± 0.02 μM for cytosolic calcium as measured by Michno *et al*., (2009) in *Drosophila* CNS neurons; 188 ± 21 μM for ER calcium as measured by Hofer *et al*., (1995) in BHK-21 cells; and 40 ± 10 nM for IP_3_ as measured by Luzzi *et al*., (1998) in *Xenopus* oocytes. While measured in various cell types that are different from those in the Drosophila pupal notum, these expected resting levels are also consistently used across many calcium signaling models (Atri et al., 1993; Li and Rinzel, 1994; Lemon et al., 2003; Han et al., 2017).

A key measure of model validity is its reproduction of wound-induced calcium signaling on similar spatial and temporal scales as observed *in vivo*. A comparison of model and experiment is shown in Figure 2, with still images at key times shown in Figure 2, A and C, and the corresponding calcium signal radius versus time graphs shown in Figure 2, A’ and C’. Similar to experiments (Movie S1), the model yields a first expansion that reaches roughly 60 μm from the wound center within ~20 s, and then recedes. It also yields an appropriate distal calcium response that becomes apparent 50-90 s after wounding and reaches cells 120-150 μm from the wound center. Finally, as in experiments, the model produces transient calcium waves or flares that extend further into the tissue. Nonetheless, the model with uniform parameters in every cell yields a calcium response that is too uniform and too radially symmetric (Figure 2B, Movie S2) despite having variations in cell size and local cell packing (Figure S1, A-C). Adding some cell-to-cell parameter variability provides a much closer match to experiments (compare Figure 2A and Figure 2C).

**Figure 2:**
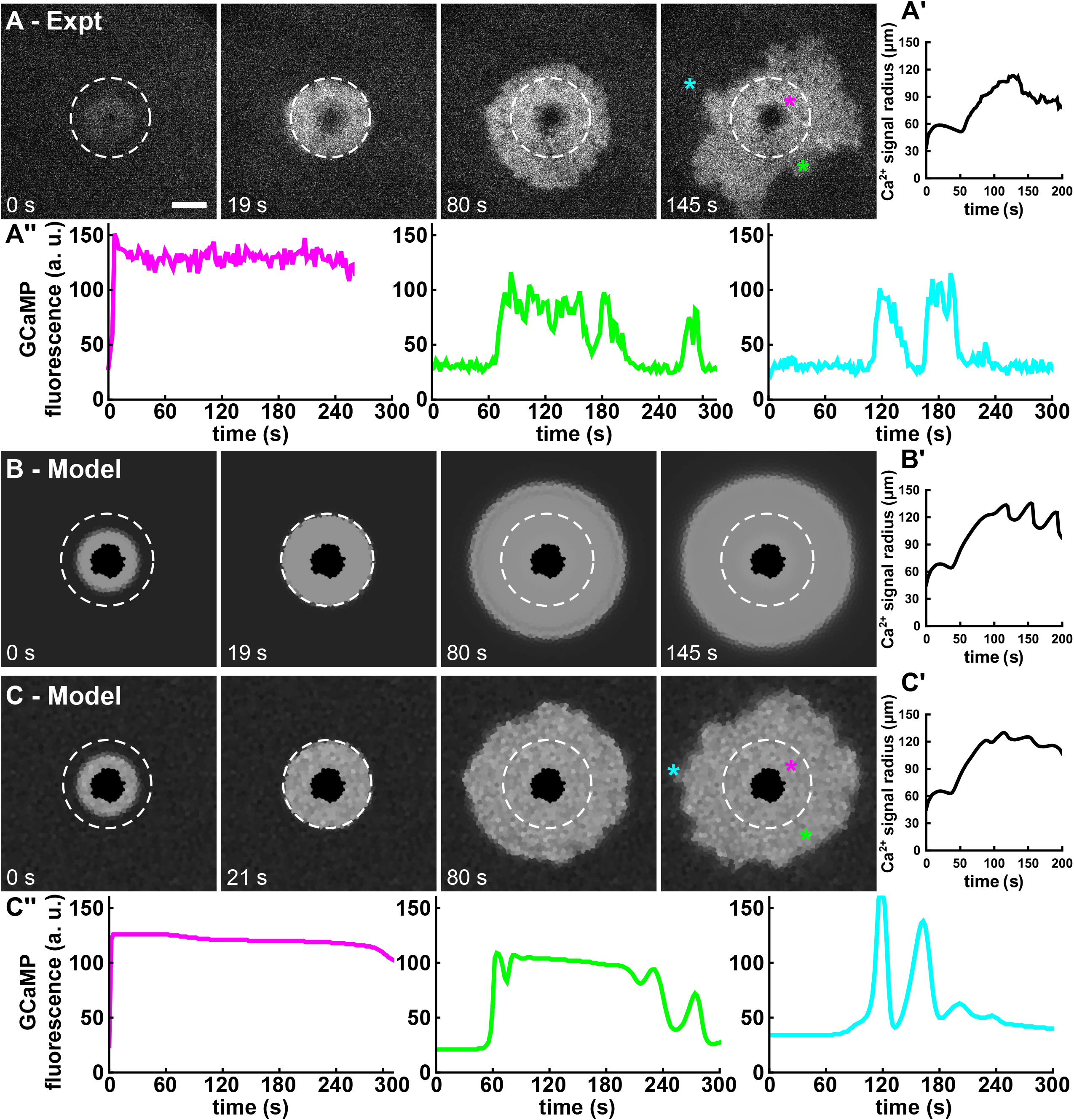
Matching the model to experiments. (A) Experimental wound-induced calcium response in control/wild-type tissue. Maximum radius of the first calcium expansion occurs at 19 s after wounding in this example and is marked by a white circle. Scale bar is 50 μm and applies to all images. (B) Model response when uniform parameters are used for all cells. (C) Model response when the IP_3_ production parameter, α, and total GCaMP parameter, B_x_, varies from cell to cell by sampling from a log-normal distribution. Such variation is needed to break radial symmetry of the calcium oscillations. (A’- C’) Corresponding quantifications of the calcium signal radius as a function of time. (A”, C”) Dynamic calcium signals at select locations that demonstrate matching single-cell dynamics in model and experiment. See also Movie S1 - Movie S3.

#### 3.1.2 The role of cellular variability

We investigated the impact of cellular variability across the tissue by running the model where a single parameter varied from cell to cell. Variability was imposed by multiplying a given base parameter from Table 2 with a cell-specific random number chosen from a log-normal distribution (associated normal distribution mean of 1 and variance of 0.2). A log-normal distribution was used to model cell-to-cell variability because it yields only positive values, and log-normal sampling has been shown to capture the impact of cell-to-cell variability on oscillatory calcium signaling (Estrada *et al*., 2016). From these single-parameter variation runs, two parameters stood out. First, varying the total GCaMP concentration within each cell, results in a minimal change in calcium signaling dynamics, but produces fluorescence variation as seen in experiments. Second, varying the IP_3_ production rate parameter, *α*, breaks the radial symmetry of the calcium waves around the wound and produces stochastic flaring behavior that matches experiments. The model output with variation in both of these parameters is shown in Figure 2, C and C’ (Movie S3) and will be taken as the base control model from here forward. Additionally, each subsequent run of the model with parameter variation utilizes a different random seed to generate a different set of randomly determined parameters for each run.

Single-parameter variability runs were performed for model parameters that do not have a set, referenced value in Table 2. These runs divide the parameters into five main groups based on the qualitative model output they produce upon variation. Variation of *α* or *η*_IPR_ produces an “ideal” response where radial symmetry is broken and calcium flares are produced. In contrast, variation of *B*_x_, *η*_GJ,Ca^2+^_, *η*_GJ,IP_3__, *J*_HC,max_, *η*_lk,PM_, *η*_μT_, or *τ*_heal_, many of which only deal with the first expansion, produces no qualitative differences in the model output as compared to the no-variation model. Variation of *w*_0_ *n_ℓ_*, or *K*_PMCA_ produces spiral waves that branch off from the region of high calcium. Spiral waves are also produced upon variation of *K*_HC_, *n*_HC_, *J*_PMCA,max_, *J*_SOC,max_, *η*_lk,ER_, *J*_SERCA,max_, or *K*_SERCA_; however, the initial conditions vary greatly from cell to cell, indicating that these parameters serve to control resting calcium levels in the model. Finally, variation of *K_ℓ_* results in calcium oscillations that arise independently of the region of high-calcium around the wound; regions of cells that are highly sensitive to the damage signal are produced in this case. Examples of these model outputs can be found in Figure S3.

#### 3.1.3 *In silico* knockdowns

As a second validation step, we performed *in silico* knockdowns and compared them to *in vivo* genetic knockdown experiments. Following methods from O’Connor et al. (2021b), experiments were run using an internally controlled experimental system: a Gal4-driven RNAi was used to knock-down a gene of interest in one section of the *Drosophila* pupal notum known as the pannier (*pnr*) domain, and a wound was made on the *pnr*-domain boundary. The wound response would thus be asymmetric, with a knockdown on one side and an internal control on the other. We generated similar *in silico* knockdowns in the model by reducing appropriate parameters for cells in just one section of the tissue mesh.

Knocking down gap junctions (*Inx2^RNAi^*) has two effects on wound-induced calcium signals (Shannon *et al*., 2017). First, because calcium can no longer move between adjacent cells, the first expansion is prevented (Figure 3, A and A’, Movie S4). In the experimental example shown, this effect is most evident via the asymmetry at 11 s after wounding. Second, the later, distal calcium response proceeds in a “speckled” pattern (Figure 3A, 120 s after wounding), i.e., the correlation between adjacent cells is greatly reduced. Nonetheless, the distal response still occurs on similar spatial and temporal scales in the knockdown and control sides of the wound (Figure 3A’). To model *in silico* knockdowns of gap junctions, we reduced the gap junction permeabilities *η*_GJ,Ca^2+^_ and *η*_GJ,IP_3__, by 90%. As shown in Figure 3, B and B’ (Movie S5), this *in silico* gap junction knockdown produces the same two characteristic impacts. Gap junction fluxes in the model thus function to allow the first expansion and coordinate large-scale correlations in the distal calcium response.

**Figure 3:**
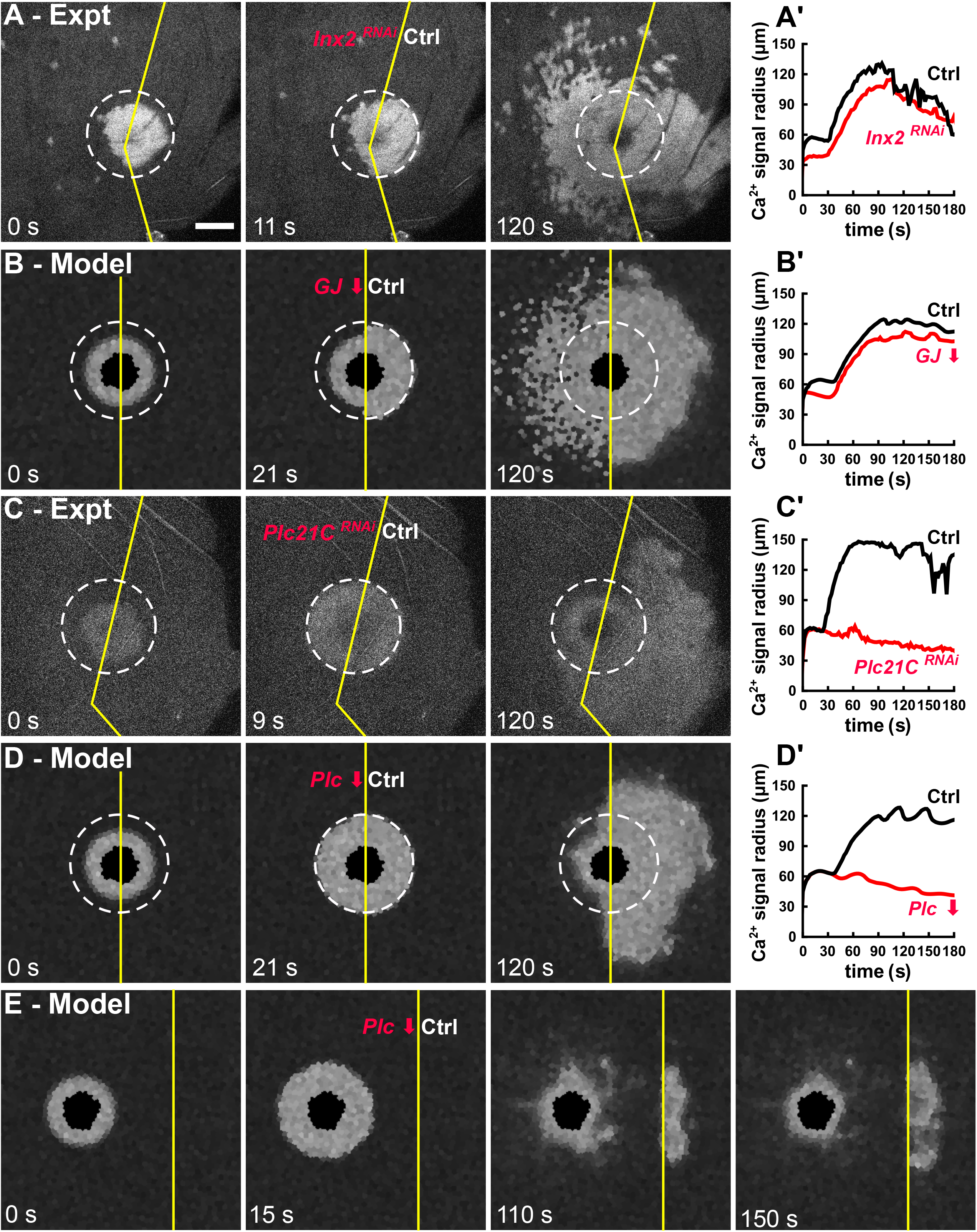
The computational model replicates knockdown experiments. (A-D) Wound-induced calcium responses with gap junctions knocked down (A-B) and PLCβ knocked down (C-D) both *in vivo* (A, C) and *in silico* (B, D). Scale bar is 50 μm and applies to all images. (A’-D’) Corresponding quantifications of the calcium signal radius on each side of the wound as a function of time. (C) Model replication of the “jump the gap” experiment; experimental data shown in O’Connor et al. (2021b). See also Movie S4Movie S8.

Knocking down any component of the G-protein cascade – *mthl10, G_αq_*, or *PLCβ* – results in a loss of the distal calcium response (O’Connor *et al*., 2021b). An experimental example of one such knockdown of *PLCβ* (*Plc21C^RNAi^*) is shown in Figure 3, C and C’ (Movie S6). The model contains a simplified version of the cascade that links the degree of Mthl10 activation to the downstream IP_3_ production rate (Eq. (24)). A reduction of the parameter *α* in Eq. (24) is thus equivalent to a *PLCβ* knockdown; it could also be interpreted as a knockdown of *G_αq_*. In either case, an *in silico* knockdown that reduces *α* by 70% (similar to RNAi-mediated knockdown efficiency (Ramos-Lewis *et al*., 2018)) leaves the first expansion unchanged, but eliminates the distal calcium response (Figure 3, D and D’, Movie S7). Direct knockdowns of the *IPR* also eliminate the distal calcium response in both experiments (O’Connor *et al*., 2021b) and the model (not shown). These observations are all strong matches to experiments.

Further, when *G_αq_* is knocked down experimentally and the tissue is wounded in the middle of the *pnr* domain rather than at the *pnr* border, the distal calcium signal appears to “jump the gap” into the control region. This experiment was performed by O’Connor et al. (2021b) to demonstrate that the signal driving the distal calcium response is propagated extracellularly without the need for any bucket-brigade mechanism. As shown in Figure 3E (Movie S8), the “jump the gap” experiment is replicated in the model by simulating a wound inside a *pnr* domain where the parameter α has been reduced by 70% (Plc↓). As expected, the distal calcium response skips over the cells with decreased α and appears at 110s after wounding only on the control side of the tissue. This match is not surprising given that the model’s reaction-diffusion equations (Eqs. (2) – (5)) explicitly include extracellular diffusion of active Gbp. Importantly, the distal signals in the control domain do not propagate back into the knockdown side in either the model or experiments. This lack of back propagation is a key check that the gap junction fluxes in the model are not too high.

Knocking down either the PMCA or the SOC does not have any noticeable effect on first/distal expansion in both experiments and model. However, these knockdowns respectively lead to higher or lower levels of calcium around the wound at later time points compared to control experiments (not shown), which is the motivation for including them in the model. Knockdowns of the SERCA pumps left many flies unhealthy and unviable for experiments, and in the model these knockdowns produce resting calcium levels way above typical physiological ranges. Therefore, the SERCA pump knockdowns were not considered for assessing the model.

### 3.2 New Insights from the Model

#### 3.2.1 Free cytosolic calcium concentrations

Interpretation of calcium signals from experimental images is limited by the properties of the fluorescent reporter used. Given GCaMP,s binding affinity for calcium (*K_x_* = 0.167 μM), its fluorescence saturates at calcium concentrations well below those typically reached during physiological signaling events (*F* – *F*_min_ ≥ 0.95(*F*_max_ – *F*_min_) when c ≥ 0.45 μM). Further, GCaMP’s high level of cooperativity (*η_x_* = 2.96) essentially gives a binary response of fluorescence “on” or “off”. Therefore, although GCaMP is useful for indicating elevated cytosolic calcium, it has limited ability to differentiate just how elevated. Although ratiometric calcium indicator dyes are better able to quantify calcium concentrations (Grynkiewicz *et al*., 1985), they cannot be used in the *Drosophila* pupal notum due to its impermeable cuticle.

The model reported here provides an alternate way to estimate the cytosolic levels of calcium around wounds. In fact, these levels are directly calculated in the model before using Eq. (30) to evaluate expected GCaMP fluorescence. Figure 4, A and B (Movie S9 and Movie S10) respectively show the GCaMP fluorescence around a control *in silico* wound and a heat map of the corresponding cytosolic calcium concentrations. The calcium concentration heat map highlights two interesting details that are not evident from GCaMP fluorescence. First, regions of rather uniform GCaMP fluorescence can hide large gradients of calcium concentration. This effect is most evident at 60 s after wounding when the calcium signals are a mix of the initial influx through plasma membrane damage, the first expansion through gap junctions, and the beginnings of IP_3_-mediated release from intracellular stores. At this time, the model reports calcium gradients between cells not adjacent to the wound as large as 1.5 μM/μm. Note that the scale of the heat map saturates at 1 μM. This choice of scale provides better visualization of distal calcium oscillations, but the calcium signals within the region of plasma membrane damage are then way off scale, reaching hundreds of μM. Second, one can observe the phasing of different parts of each calcium wave or flare. For example, there is an oscillation just trailing the front of the distal expansion that is not evident in the fluorescence signals. The model calculations provide a cautionary example that cells within the calcium signaling region can have similar GCaMP fluorescence levels and yet experience drastically different calcium concentration histories.

**Figure 4:**
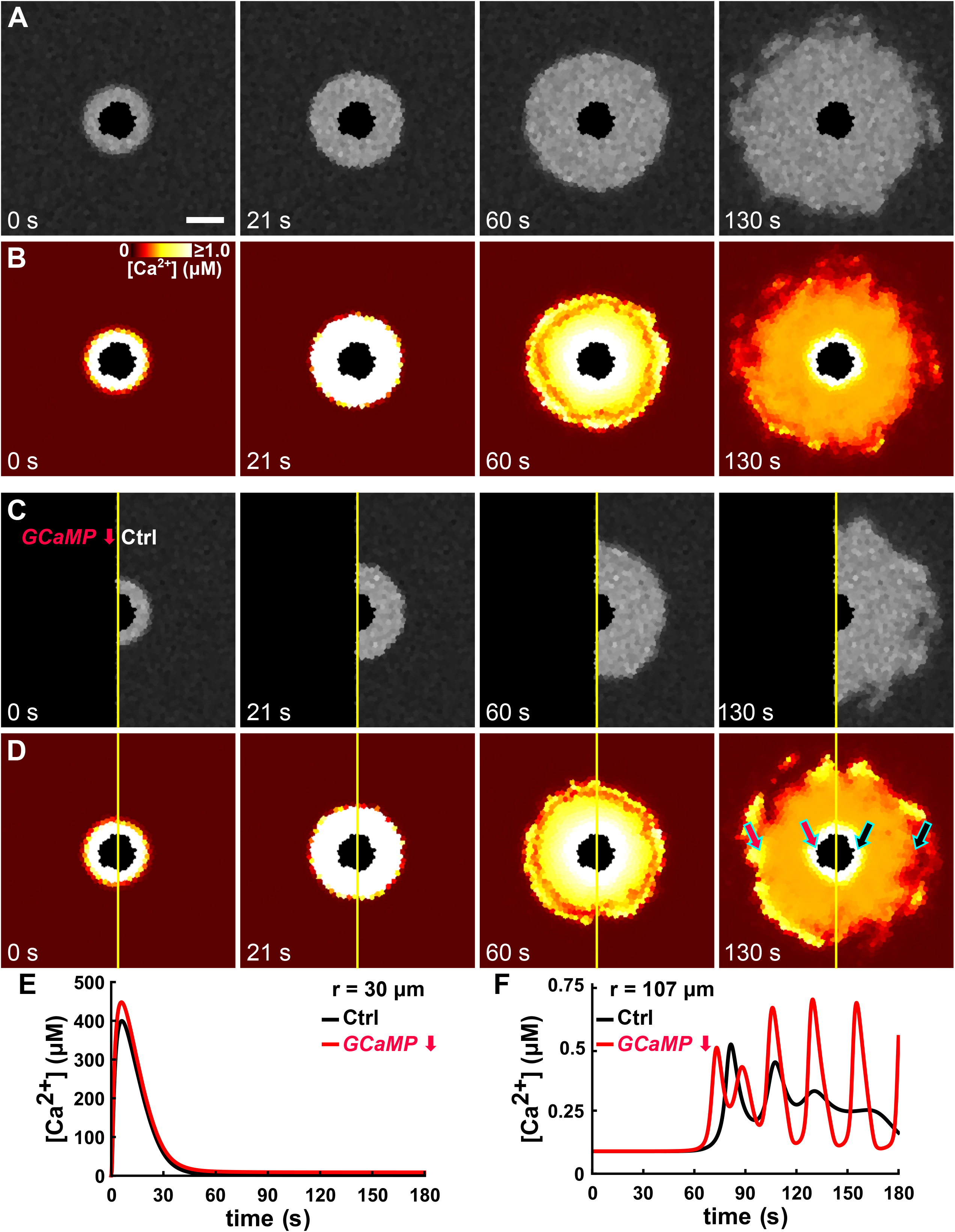
The model elucidates hidden structure in the free calcium concentrations. (A-B) GCaMP fluorescence and corresponding free cytosolic calcium concentration in a control system. (C-D) Same in a GCaMP knockdown. Scale bar is 50 μm and applies to all images. The color scale for free cytosolic calcium is shown in B and applies to all images in B and D. Note that the scale saturates at 1.0 μM and that the wound region has been colored black since it is not cytosolic. (C) Free cytosolic calcium versus time for cells close to the wound; locations as marked by inner arrows in D. (F) Same for cells further from the wound; locations marked by outer arrows in D. See also Movie S9Movie S12.

#### 3.2.2 GCaMP’s effect on calcium signals

The fluorescent calcium reporter GCaMP is also a buffer of calcium. The ability of buffers to impact calcium oscillations has been studied extensively both experimentally and theoretically; buffers decrease oscillation amplitude and frequency, even to the point of preventing calcium oscillations and waves altogether (Nuccitelli *et al*., 1993; Wagner and Keizer, 1994; Skupin *et al*., 2008). Given how information in calcium signals can be encoded by both amplitude and frequency (Thomas *et al*., 1996; Berridge *et al*., 1998; Brodskiy *et al*., 2019), it is important to understand how the presence of GCaMP alters calcium signals following wounding.

We therefore ran an *in silico*, internally controlled knockout of GCaMP. Experimentally, this would be useless: no GCaMP would mean no fluorescent reporter of calcium; however, as shown in Figure 4, C and D (Movie S11 and Movie S12), the model can directly report the free cytosolic calcium concentration, with or without GCaMP. The absence of GCaMP as a buffer has three modest but discernable effects. First, the initial influx yields a ~10% higher peak of free cytosolic calcium levels (Figure 4E). Second, the distal calcium expansion occurs slightly earlier (Figure 4F). Third, the absence of GCaMP leads to distal calcium oscillations with a larger amplitude and frequency (Figure 4F). Consistent with observations of GCaMP expression not disrupting biological processes (Chen *et al*., 2013), the size of these impacts is small under the levels of GCaMP used here, but they would increase if GCaMP were expressed more strongly.

#### 3.2.3 The roles of intercellular calcium and IP_3_ transfer

Intercellular coupling of cells is a necessary condition for intercellular calcium waves to initiate and persist across a tissue (Leybaert and Sanderson, 2012). The model presented here allows for the gap-junction transfer of both cytosolic calcium and IP_3_ (Eqs. (27) and (28)). Although knockdown experiments cannot individually determine the importance of each species’ gap junction flux, the model presented here and matched to experiments does provide such insight.

During the first expansion, the amount of calcium moving through gap junctions is ~1000-fold higher than IP_3_ (Figure 5A); however, during the distal expansion and subsequent flares, calcium and IP_3_ have similarly sized fluxes (Figure 5B). This difference suggests that the relative importance of each flux may change with time after wounding. Nonetheless, the relative fluxes do not necessarily equate to importance. One must also consider how the fluxes change each species’ concentration and how those concentrations compare to the half-maximal effect levels for downstream steps.

**Figure 5:**
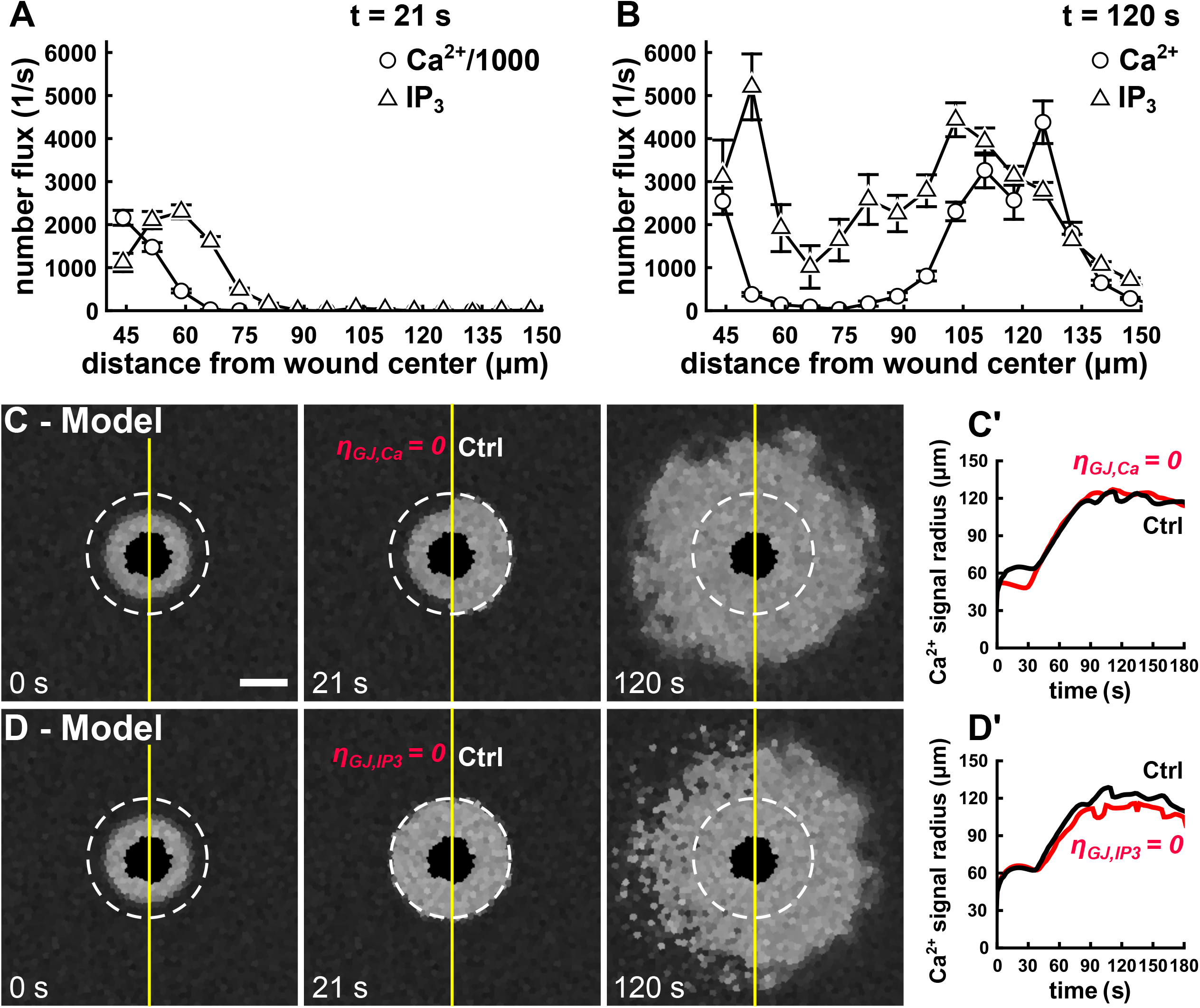
Model demonstrates different roles for gap-junction fluxes of calcium and IP_3_. (A-B) Quantification of average gap-junction fluxes, binned by distance from the wound center at both the maximum extent of the first expansion (21 s after wounding) and just after completion of the second distal expansion (120 s). Error bars denote standard error of the mean. Note that the calcium fluxes in A have been divided by a factor of 1000 to plot on the same scale as IP_3_ fluxes. (C-D) Model responses after selective *in silico* knockdown of gapjunction fluxes for either calcium (C) or IP_3_ (D). Scale bar is 50 μm and applies to all images. (C’-D’) Corresponding quantifications of the calcium signal radius as a function of time. See also Movie S13 and Movie S14.

The model does provide a path to exploration currently unavailable in experiments: we can run simulations in which one species’ gap junction flux is eliminated but the other is left unchanged. To highlight the impact of each flux elimination, we run the simulations as an internally controlled knockdown system. When only the calcium flux is eliminated, the first expansion is nonexistent (Figure 5C, Movie S13; *η*_GJ,Ca_ = 0). This result matches the effect of a full gap junction knockdown on the first expansion (Figure 3, A and B, *t* = 11 or 21s). On the other hand, eliminating the calcium flux still allows for a coordinated distal expansion and flares. This result does not match the “speckled” pattern evident with a full gap junction knockdown (Figure 3, A and B, *t* = 120 s). This speckled pattern can be reproduced if the gap junction fluxes are altered so that IP_3_ fluxes are eliminated (Figure 5D, Movie S14; *η*_GJ,IP_3__ = 0). Setting both gap junction permeabilities to zero produces both effects (data not shown), similar to the 90% gap-junction knockdown shown in Figure 3C. These model results predict that intercellular transfer of calcium is necessary for the first expansion, while intercellular transfer of IP_3_ coordinates multicellular calcium signaling during the later distal expansion and flares.

## 4 Discussion

We have developed a mathematical model to replicate and predict calcium signaling events following laser-induced wounding of the *Drosophila* pupal notum. At the tissue level, the model incorporates both a simple model of physical damage and a reaction-diffusion model of biochemical damage signals. These damage models are applied across the field of cells which share a common calcium signaling toolkit (Figure 1). The calcium signaling toolkit elements and their associated parameters (Table 2) were chosen to replicate the spatiotemporal dynamics of calcium signaling events that follow wounding of wild-type tissues (Figure 2) and genetic knockdowns (Figure 3).

The model was then used to investigate the system in ways that are not currently experimentally accessible. First, the model reveals that many of the calcium signals are well above the K_d_ of the GCaMP sensor, and thus regions with rather uniform fluorescence can hide order-of-magnitude differences in cytosolic calcium concentration (Figure 4): cells within the first expansion experienced peak calcium levels of 100-400 μM, but cells within the distal expansion reached peaks around just 0.5 μM. Second, by running the model with or without the genetically-encoded calcium indicator GCaMP, we could show that GCaMP buffering has a small effect on the frequency and amplitude of calcium oscillations or flares, but has little to no effect on the spatial and temporal scales of the first and distal expansion. Third, by modeling selective gap junction knockdowns, i.e., allowing only calcium or IP_3_ but not both to move between adjacent cells, the model shows that gap-junction fluxes of calcium play a key role in the first expansion, but it is the gap-junction transfer of IP_3_ that couples and coordinates cells during the distal expansion and subsequent flares (Figure 5).

### 4.1 Cell-to-cell variability

Cell-to-cell variability across the tissue is a necessary condition for the model to break radial symmetry in the calcium flares (Figure 2). We find that variations in two parameters could yield the appropriate broken symmetry without having other adverse effects: α, the parameter that controls the maximum IP_3_ production rate (Eq. (24)); or ηIPR, the permeability of the fully open IPR (Eq. (18)). These two parameters respectively control how much IP_3_ will be produced for a given level of damage signal Gbp and how much calcium is subsequently released from the ER, and thus they control the behavior of the IP_3_-dependent calcium flares (Figure 3, C and D). Both parameters also had minimal impact on the resting calcium levels of modeled cells. We decided to move forward with variations in α because it represents the activity of an entire G-protein cascade, and the output of this multistep cascade would include the variability generated within each step of this signaling network, thus resulting in a multiplicative version of the central limit theorem where the result approaches a lognormal distribution (Limpert *et al*., 2001).

The necessity of cell-to-cell variability begs a key question: is such variability simply noise or is it a key aspect of wound-induced calcium signaling? Analyses of other systems have shown that cell-to-cell variability is not intrinsic noise that obscures a signal, but instead increases the capacity of a group of cells to transmit information (Wada *et al*., 2021). Further, as Yao et al. (2016) demonstrated, variations in cellular calcium responses can be explained by structured variability in cell state. Our model is thus far agnostic as to the role of cell-to-cell variability, but future investigations that link wound-induced calcium signals to downstream cellular behaviors should strive to discern whether the clearly present variability serves a functional role (Yao *et al*., 2016).

### 4.2 Role of intercellular calcium and IP_3_ transfer

In the model, both calcium and IP_3_ travel from a cell to its neighbors through gap junctions. By modeling “selective” gap junctions where only calcium or IP_3_ is allowed to move intercellular, we were able to tease apart separate roles for both fluxes. Given that the first expansion is caused by large amounts of calcium rushing in through plasma membrane damage, it is not surprising that intercellular calcium flux is necessary for the first expansion. Smaller calcium fluxes are still present during the distal calcium response and flares, but these fluxes are not sufficient to locally coordinate calcium responses. Instead, coordination of the distal expansion and flares is primarily driven by the gap-junction transfer of IP_3_ (Figure 5, C and D) – a result consistent with many other systems (Leybaert and Sanderson, 2012). The selective gap junction simulations thus piece together the full gap junction knockdown experiments and simulations (Figure 3, A and B): the first expansion is prevented due to a lack of calcium transfer, and the distal calcium response is altered due to a loss of IP_3_ transfer.

Interestingly, the different roles for calcium and IP_3_ transfer occur at later times in the simulations even though the radial fluxes of both species are similar. Obviously, the impacts of those fluxes are not equivalent. One reason is that cytosolic calcium concentrations are strongly buffered (Eq. (8)). This buffering reduces the impact of calcium fluxes unless they are large enough to overwhelm the cells buffering capacity, as is the case during the earlier first expansion. IP_3_ has no such buffering present. In addition, small changes in IP_3_ levels are leveraged into larger changes in cytosolic calcium through the Hill functions that link IP_3_ to the opening of IPR and release of calcium from ER stores (Eqs. (18) and (19)).

### 4.3 Comparison to other calcium signaling models

The model developed here goes well beyond currently published models of wound-induced calcium signaling. Narciso et al. (2015) developed a calcium signaling model in the context of laser-ablation wounding of *Drosophila* wing discs; however, their wound is modeled only implicitly as a time-integrated IP_3_ stimulus over a small circular area.

Additionally, their model is implemented over a two-dimensional, homogenized continuum rather than over discrete cells in a tissue. A more recent model from the same lab investigates calcium signaling across a *Drosophila* wing disc, but with a focus on organ development and with the total IP_3_ production rate in each cell being just a function of the cytosolic calcium concentration (Soundarrajan *et al*., 2021).

Donati et al. (2021) developed a model of calcium signals in mouse keratinocytes upon photodamage; however, instead of proteolytic activation driving the extracellular signaling mechanism, the damage signal in that system is ATP, which is released both from the wound site as well as from undamaged cells through connexin hemichannels. O’Connor et al (2021b) found that the adenosine receptor played no role in the wound-induced calcium response in the *Drosophila* notum.

Here we develop a more detailed model that can be used to describe calcium signaling around an explicit wound with both physical and biochemical damage signals. Further, our model is based on experiments in *Drosophila*. Although our chosen model components are not unique to *Drosophila*, a model that captures calcium signaling in this system is advantageous due to its utility. *Drosophila* has a well-established genetic toolkit, which provides an accessible route to discerning specific molecular mechanisms. Experimental genetic manipulations were key to outlining the components needed in the model and provide a stringent test for evaluating the model’s *in silico* knockdowns.

Continuing studies that tightly align expansions of the model presented here with future genetic experiments should provide further insights linking calcium signaling to wound healing. A promising future direction is to relate specific calcium signaling events to downstream wound healing responses such as cytoskeletal remodeling, actomyosin purse string formation, cell migration, and reepithelization. Multiple groups have used computational modeling to investigate cell and tissue mechanics around epithelial wounds (Staddon *et al*., 2018; Ajeti *et al*., 2019), feedback between cell mechanics and calcium (Kaouri *et al*., 2019), and decoding mechanisms that initiate calcium-dependent transcription factors (Noren *et al*., 2016; Brodskiy and Zartman, 2018). It should be possible to link the model presented here to these downstream response models, but with a key modification: the model presented here considers relatively short time scales of seconds to minutes after wounding, for which cell and tissue geometry is reasonably static, but the downstream response models consider longer time scales of tens of minutes to hours where wound healing clearly requires dynamic geometries. The challenge will be adapting the current model to function across cells that move, change shape, and exchange neighbors. Doing so will be a key step in linking to existing wound response models and relating wound-induced calcium signals to downstream wound healing mechanisms.

### 4.4 Conclusion

Laser-wounds in the *Drosophila* pupal notum give rise to a complex set of calcium signaling events. These events are driven by at least two different aspects of the wound, and yet interact with a common set of cellular components to generate overlapping calcium signals. Here, we have developed a mathematical model that includes both physical and chemical damage signals and integrates them with a single calcium signaling toolkit at the cellular level. The model replicates experimental results in wild-type and genetically manipulated tissues and yields key insights into cell-to-cell variability, cytosolic free calcium behavior, and the roles of intercellular gap-junction communication. It provides an important framework for future studies of both calcium signaling and wound responses in the *Drosophila* system.

## Supporting information

Main figures with captions

## Abbreviations used

ER: endoplasmic reticulum;
Gbp: growth-blocking peptide;
GCaMP: a genetically-encoded calcium indicator,
GJ: gap junction;
Gαq: α subunit of a G-protein;
HC: high capacity;
IP_3_: inositol trisphosphate;
IPR: IP_3_ receptor;
Mthl10: Methuselah-like 10;
ODE: ordinary differential equation;
PDE: partial differential equation;
PLC: phospholipase C;
PM: plasma membrane;
PMCA: plasma membrane calcium ATPase;
SERCA: sarcoendoplasmic reticulum calcium ATPase;
SOC: store-operated calcium channel;
μT: micro-tear

## 5 Acknowledgements

This work was supported by the National Institute of General Medical Sciences (1R01GM130130 to A.P.-M. and M.S.H.). J.T.O. was supported by the National Institute of Child Health and Human Development (T32HD007502) and the American Heart Association (19PRE34410069 to J.T.O.). This work was conducted in part using the resources of the Advanced Computing Center for Research and Education (ACCRE) at Vanderbilt University, Nashville, TN. We thank the Bloomington Drosophila Stock Center (Bloomington, IN) and the National Institute of Genetics (Shizuoka, Japan) for Drosophila stocks and G. Struhl (Columbia University, New York, NY) for providing the plasmid containing the Actin5c promoter.

