## Supplemental Information for "A mathematical model of calcium signals around laser-induced epithelial wounds"

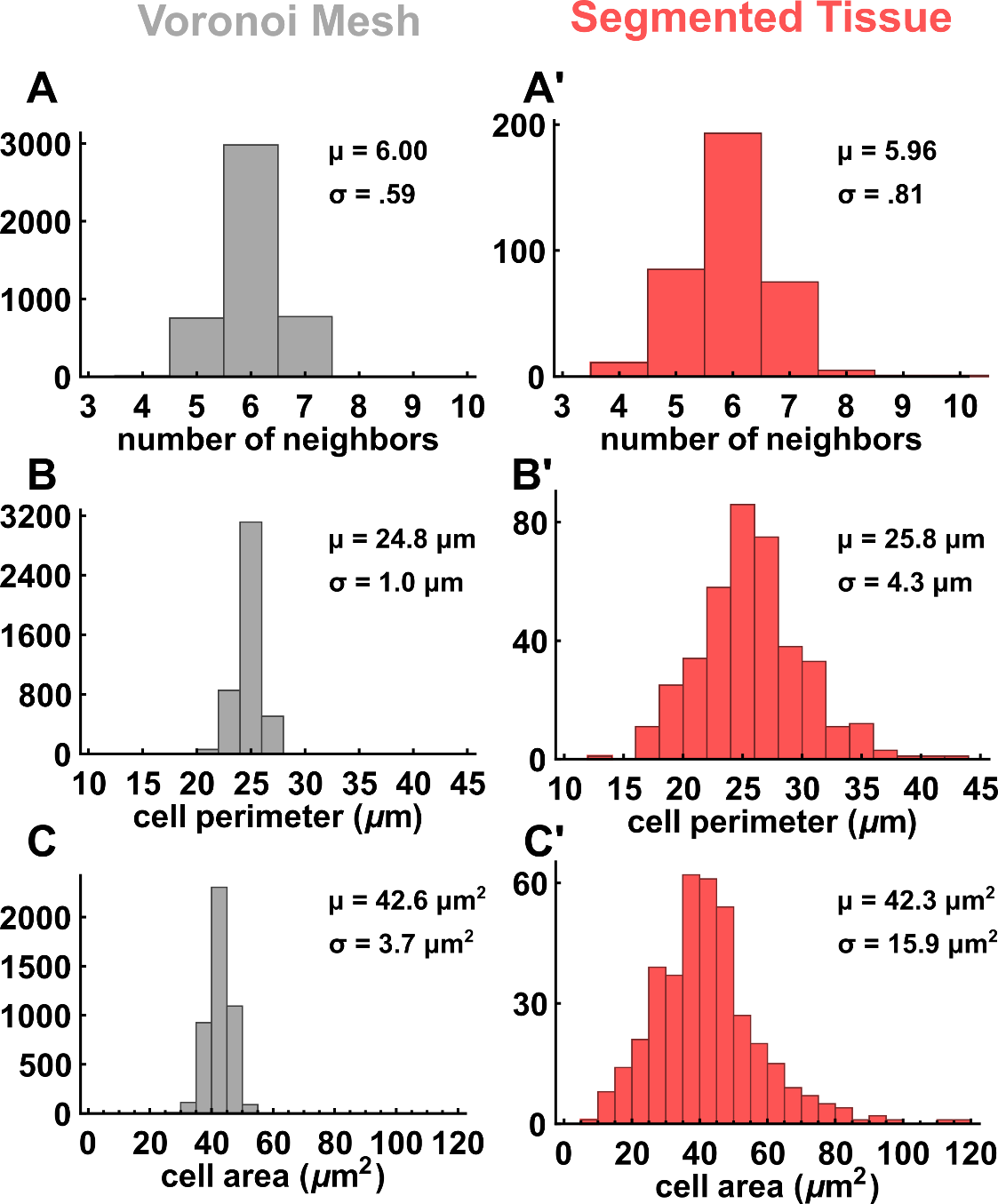

Figure S1: Cell packing and geometry distributions for the Voronoi mesh and a segmented tissue

Distributions for number of neighbors (A, A’), cell perimeter (B, B’) and cell area (C, C’) for the Voronoi mesh (A – C) and a segmented 163 μm x 163 μm section of an *in vivo* tissue (n = 390 cells) (A’ – C’). Means (μ) and standard deviations (σ) are reported for each distribution.

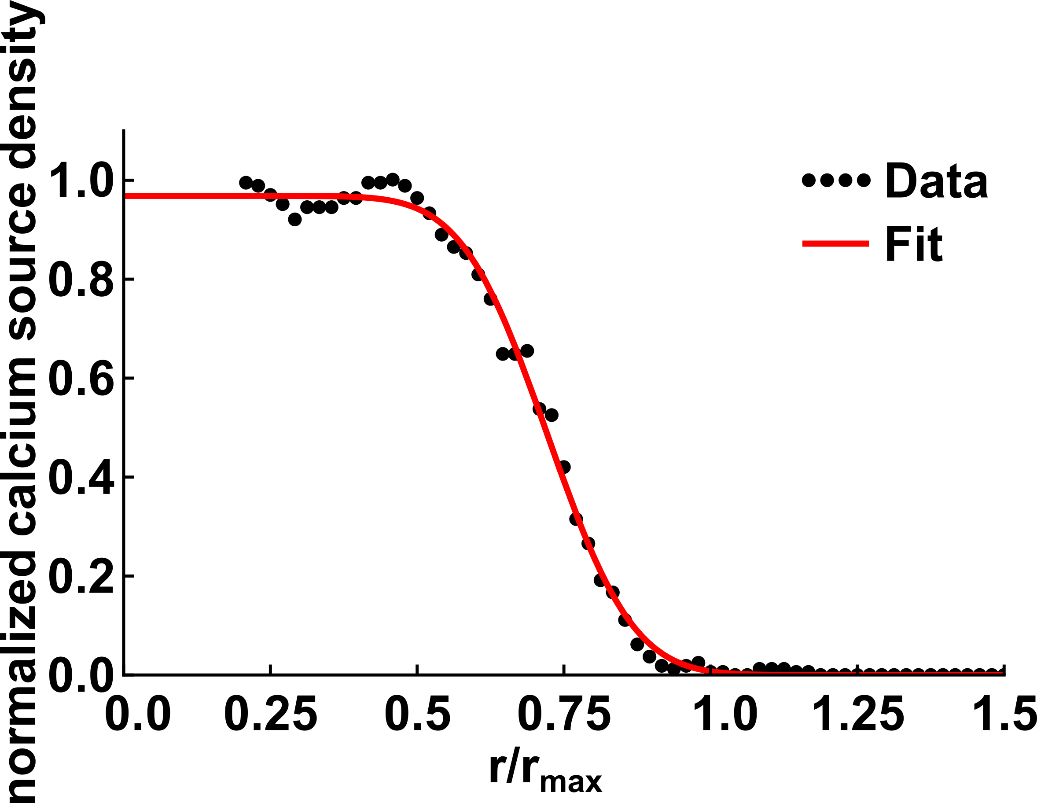

Figure S2: Calcium source density is fit to a heuristic function.

Image analysis of GCaMP fluorescence 17 ms after wounding yields a radial density of calcium entry sites as a function of distance from the wound (black points). The data is fit to Eq. (1) (red line). The vertical axis is normalized so that the maximum density is 1, and the horizontal axis is scaled relative to the maximum radius of the cavitation bubble.

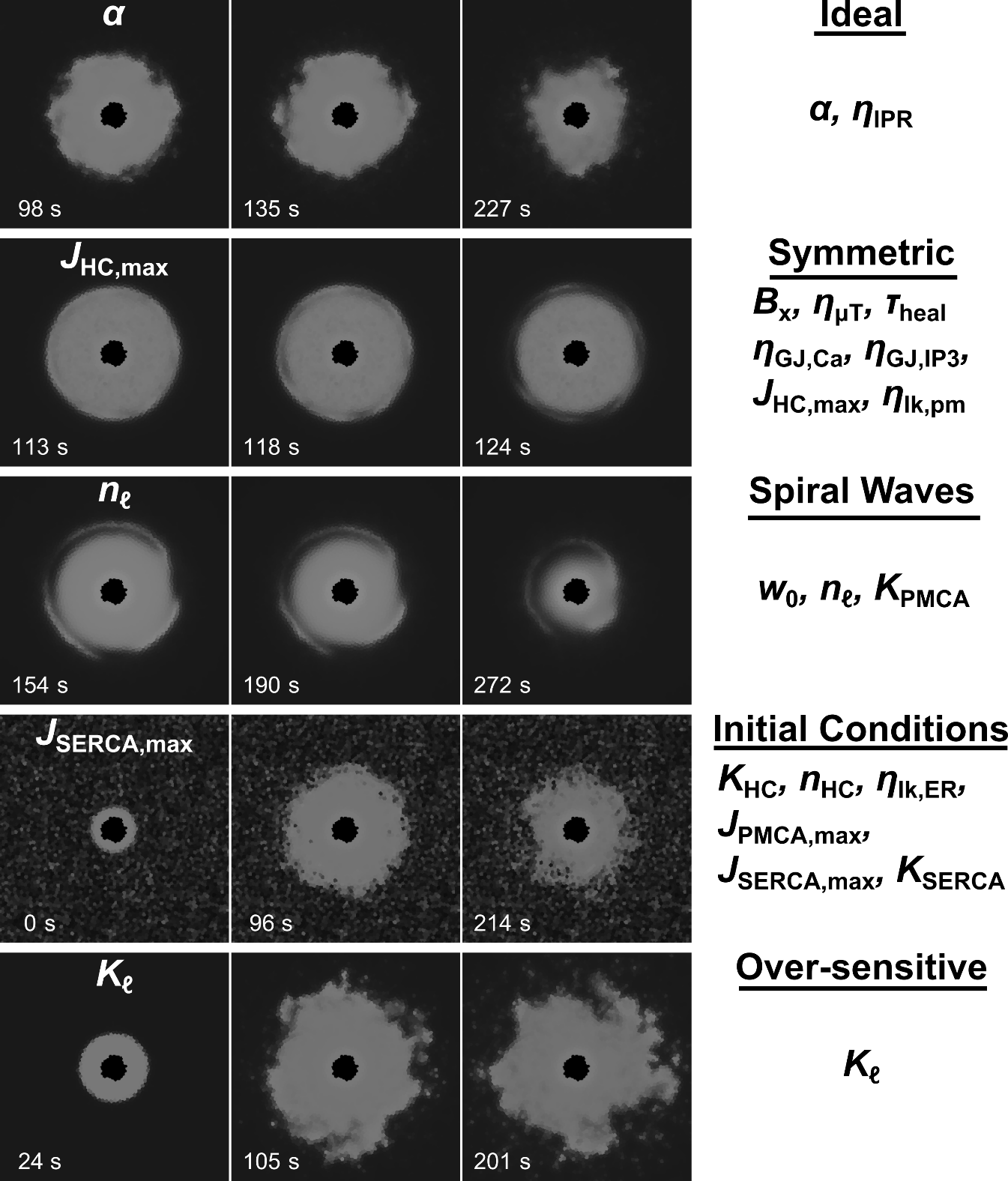

Figure S3: Example single-parameter variation model outputs.

Parameters are grouped based on the model output when only that parameter is varied across the tissue. One example output of each group is shown in each row, with the full list of parameters in each group in the right-most column. Frames were chosen that best show the group behavior. “Ideal”: model outputs where radial symmetry is broken and calcium flares are produced. “Symmetric”: model outputs where calcium oscillations are radially symmetric, similar to having no parameter variation. “Spiral Waves”: Model outputs where spiral waves emerge around the high-calcium region. “Initial Conditions”: model outputs where initial conditions have large variations; spiral waves are also present. “Over-sensitive”: model output where flares and calcium oscillations arise independently of the high-calcium region.

**Supplemental movie legends**

Movie S1, related to Figure 2A: Experimental wound-induced calcium response in control/wild-type tissue

Movie S2, related to Figure 2B: Model response when uniform parameters are used for all cells

Movie S3, related to Figure 2C: Model response when the IP­_3_ production parameter, α, and the total GCaMP parameter, B_x_, varies from cell to cell by sampling from a log-normal distribution. Such variation is needed to break radial symmetry of the calcium oscillations.

Movie S4, related to Figure 3A: Experimental wound-induced calcium response with gap junctions knocked down (left side) in an internally controlled tissue

Movie S5, related to Figure 3B: Model wound-induced calcium response with gap junctions knocked down (left side) in an internally controlled tissue

Movie S6, related to Figure 3C: Experimental wound-induced calcium response with PLCβ knocked down (left side) in an internally controlled tissue

Movie S7, related to Figure 3D: Model wound-induced calcium response with PLCβ knocked down (left side) in an internally controlled tissue

Movie S8, related to Figure 3E: Model replication of the “jump the gap” experiment; experimental data shown O’Connor et al. (2021b).

Movie S9, related to Figure 4A: GCaMP fluorescence in a control model system corresponding to the free cytosolic calcium concentrations in Movie S10.

Movie S10, related to Figure 4B: Free cytosolic calcium in a control model system. Scale bar shown in Figure 4B

Movie S11, related to Figure 4C: GCaMP fluorescence in an internally controlled GCaMP knockdown system (left side) corresponding to the free cytosolic calcium concentrations in Movie S12

Movie S12, related to Figure 4D: Free cytosolic calcium concentration in an internally controlled GCaMP knockdown system (left side). Scale bar shown in Figure 4B.

Movie S13, related to Figure 5C: Model wound-induced calcium response without intercellular calcium transfer (left side) in an internally controlled tissue

Movie S14, related to Figure 5D: Model wound-induced calcium response without intercellular IP_3_ transfer (left side) in an internally controlled tissue
