## Supplementary material for "A mathematical model of calcium signals around laser-induced epithelial wounds": Main figures with captions

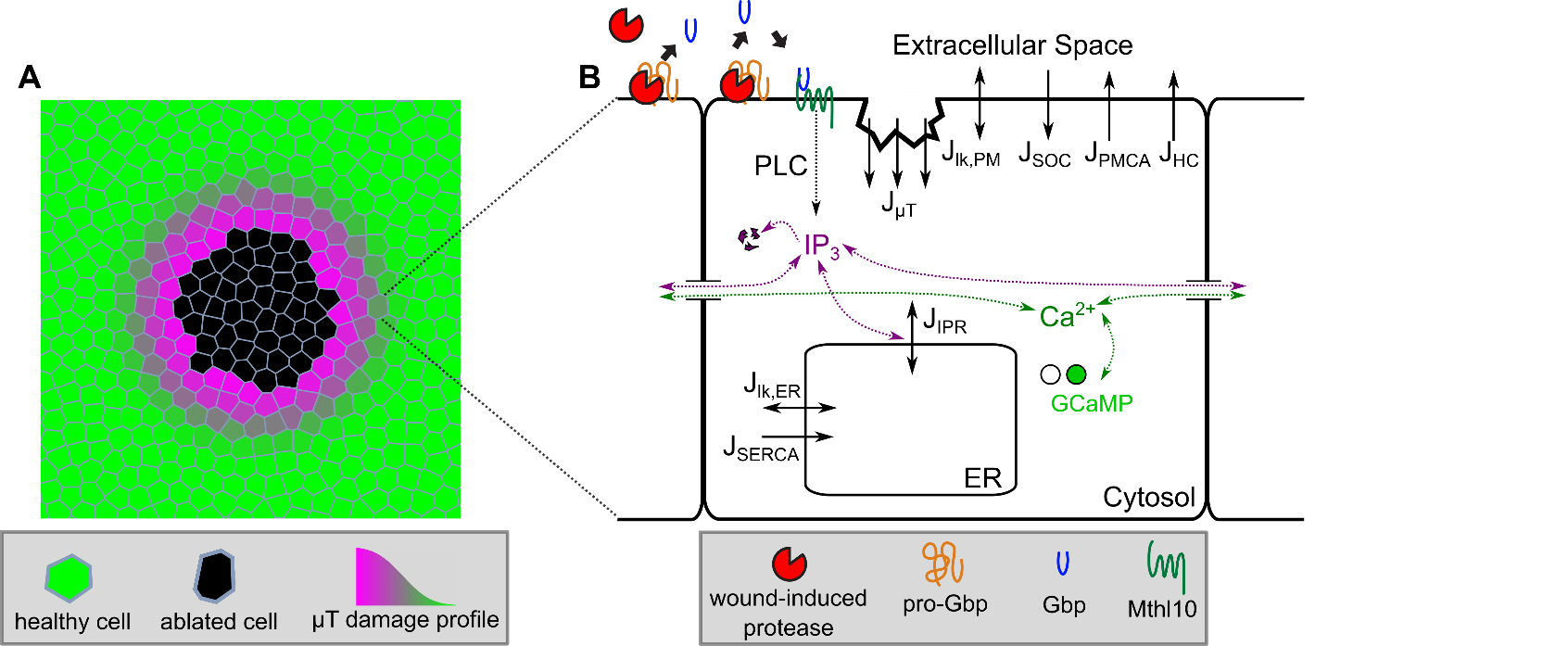

Figure 1: Schematic diagram of the complete wound-induced calcium signaling model

160 x 160 μm section of the tissue mesh centered around the wound. Cells are color-coded according to their physical damage level. (B) Reactions and fluxes included in the single-cell model and its connection to the extracellular model of biochemical damage signals. Solid black arrows show directions of calcium fluxes.

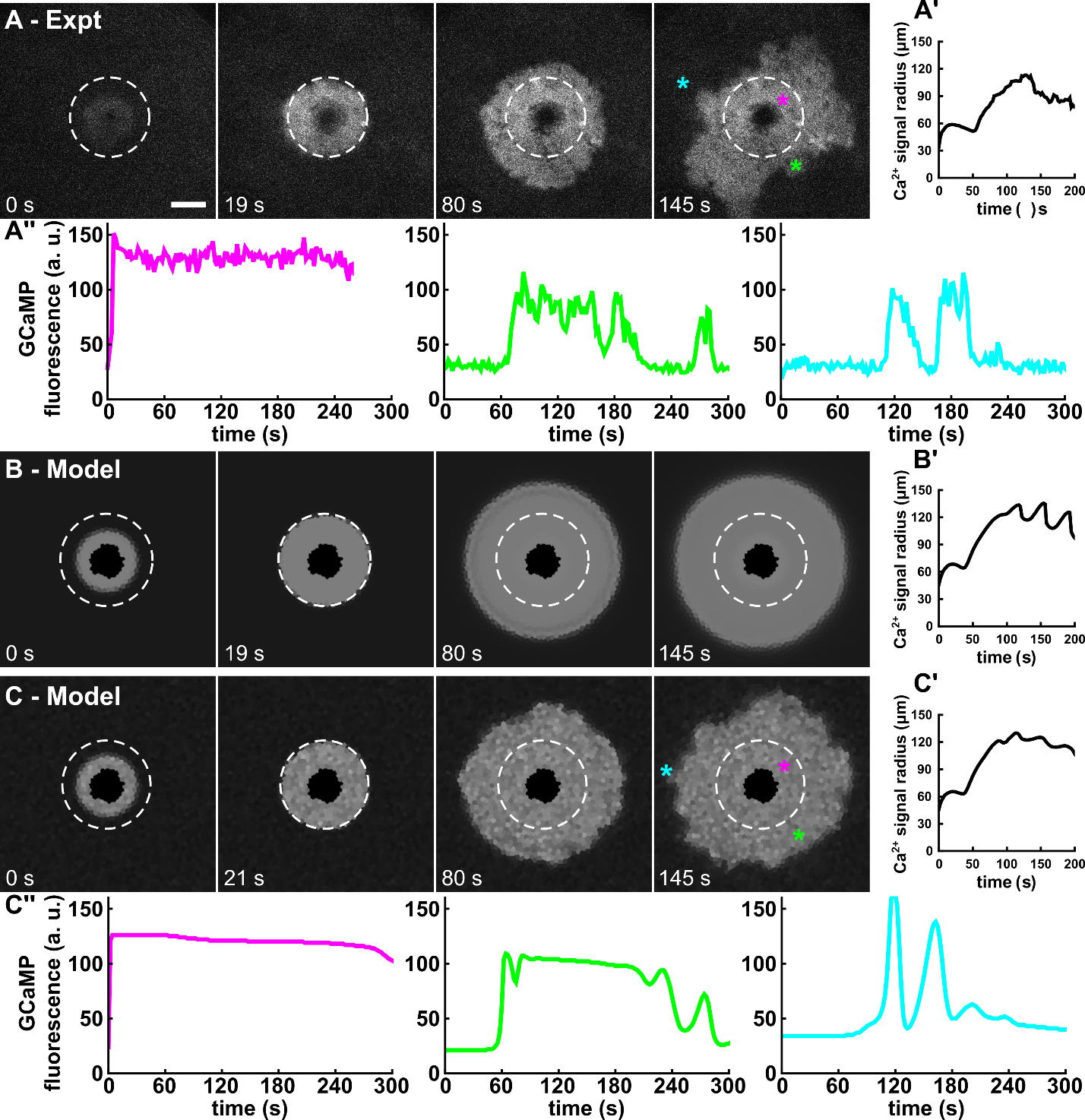

Figure 2: Matching the model to experiments

(A) Experimental wound-induced calcium response in control/wild-type tissue. Maximum radius of the first calcium expansion occurs at 19 s after wounding in this example and is marked by a white circle. Scale bar is 50 μm and applies to all images. (B) Model response when uniform parameters are used for all cells. (C) Model response when the IP­_3_ production parameter, α, and total GCaMP parameter, B_x_, varies from cell to cell by sampling from a log-normal distribution. Such variation is needed to break radial symmetry of the calcium oscillations. (A’-C’) Corresponding quantifications of the calcium signal radius as a function of time. (A’’, C’’) Dynamic calcium signals at select locations that demonstrate matching single-cell dynamics in model and experiment. See also Movie S1 - Movie S3.

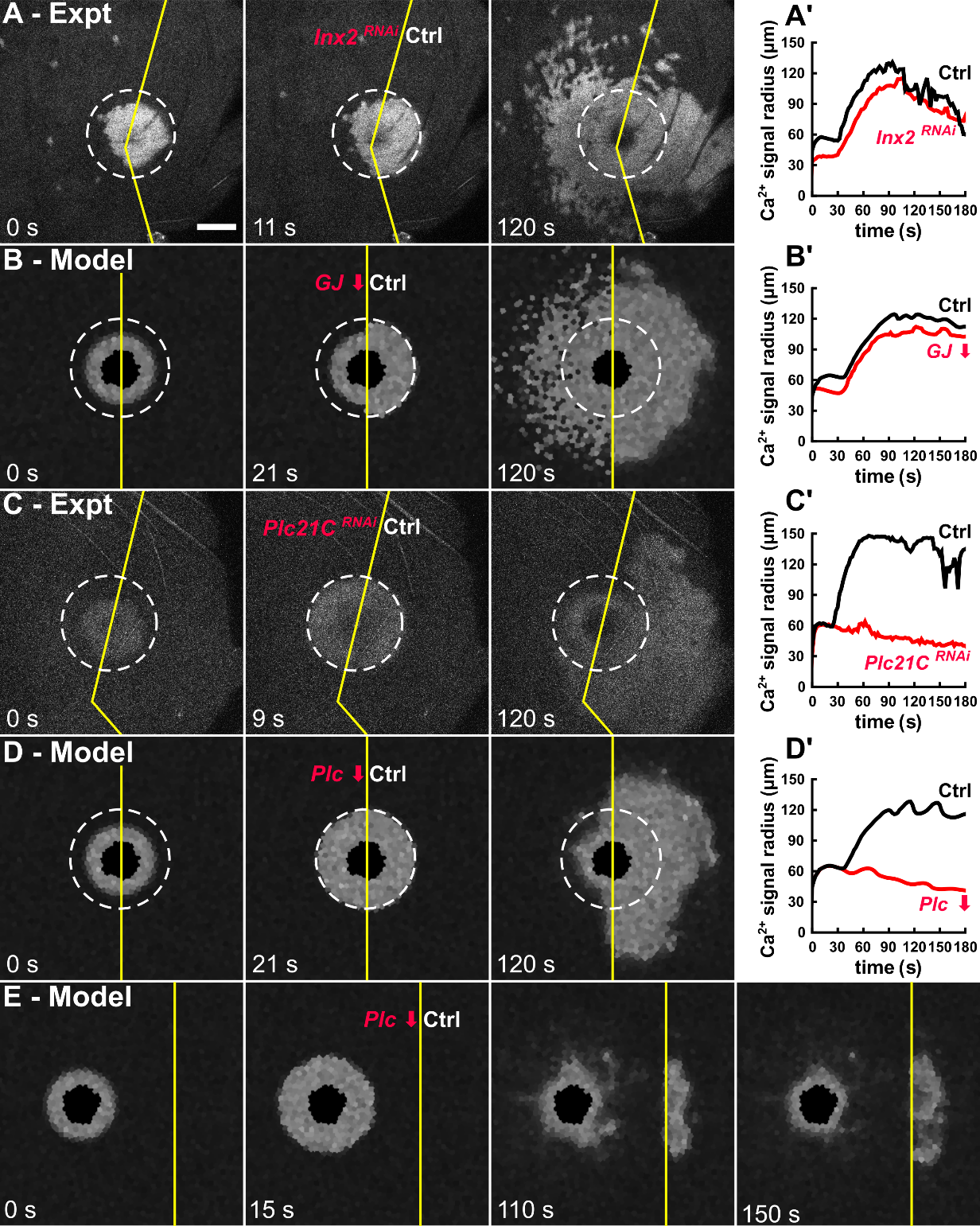

Figure 3: The computational model replicates knockdown experiments.

(A-D) Wound-induced calcium responses with gap junctions knocked down (A-B) and PLCβ knocked down (C-D) both *in vivo* (A, C) and *in silico* (B, D). Scale bar is 50 μm and applies to all images. (A’-D’) Corresponding quantifications of the calcium signal radius on each side of the wound as a function of time. (C) Model replication of the “jump the gap” experiment; experimental data shown in O’Connor et al. (2021b). See also Movie S4Movie S8.

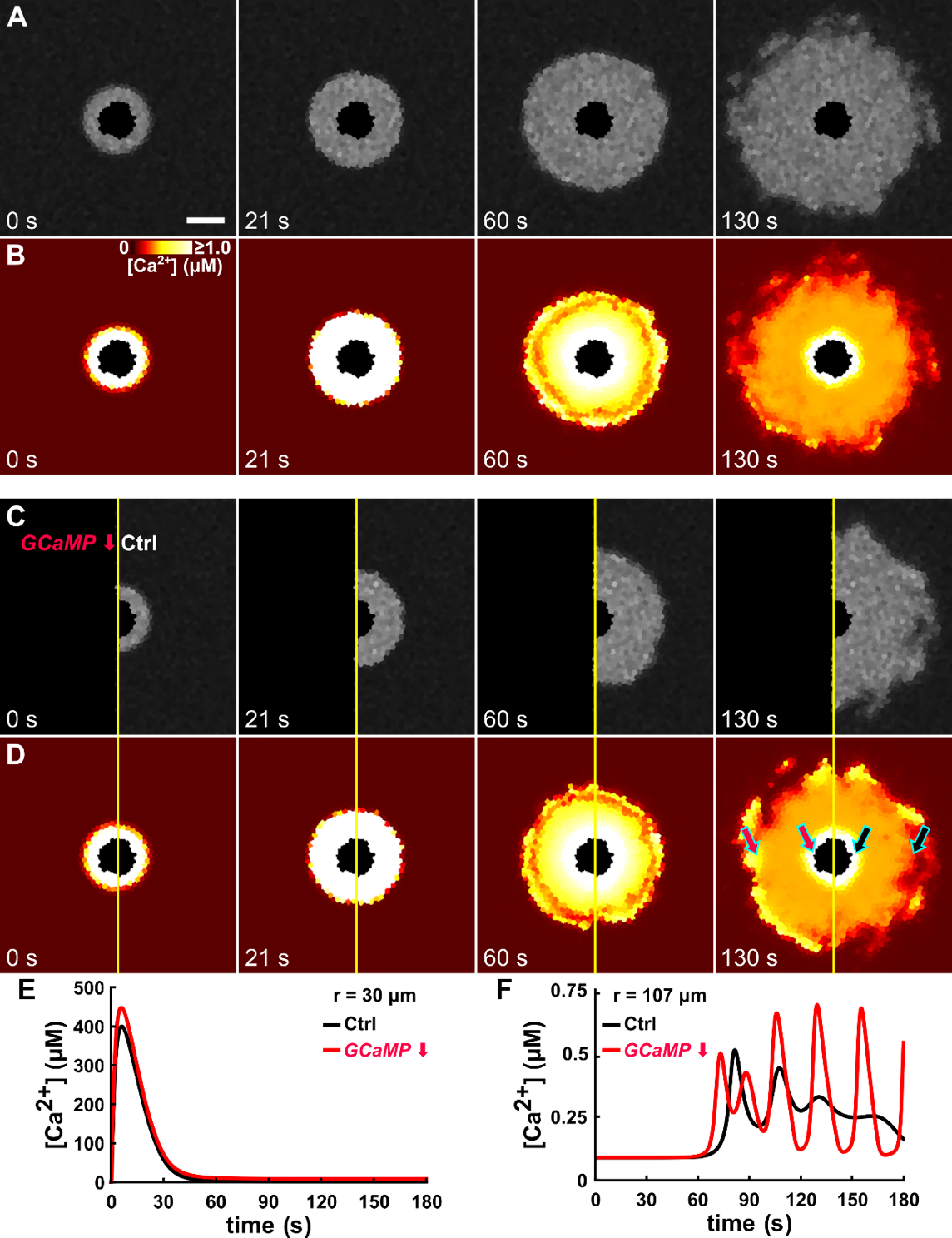

Figure 4: The model elucidates hidden structure in the free calcium concentrations.

(A-B) GCaMP fluorescence and corresponding free cytosolic calcium concentration in a control system. (C-D) Same in a GCaMP knockdown. Scale bar is 50 μm and applies to all images. The color scale for free cytosolic calcium is shown in B and applies to all images in B and D. Note that the scale saturates at 1.0 μM and that the wound region has been colored black since it is not cytosolic. (C) Free cytosolic calcium versus time for cells close to the wound; locations as marked by inner arrows in D. (F) Same for cells further from the wound; locations marked by outer arrows in D. See also Movie S9 - Movie S12.

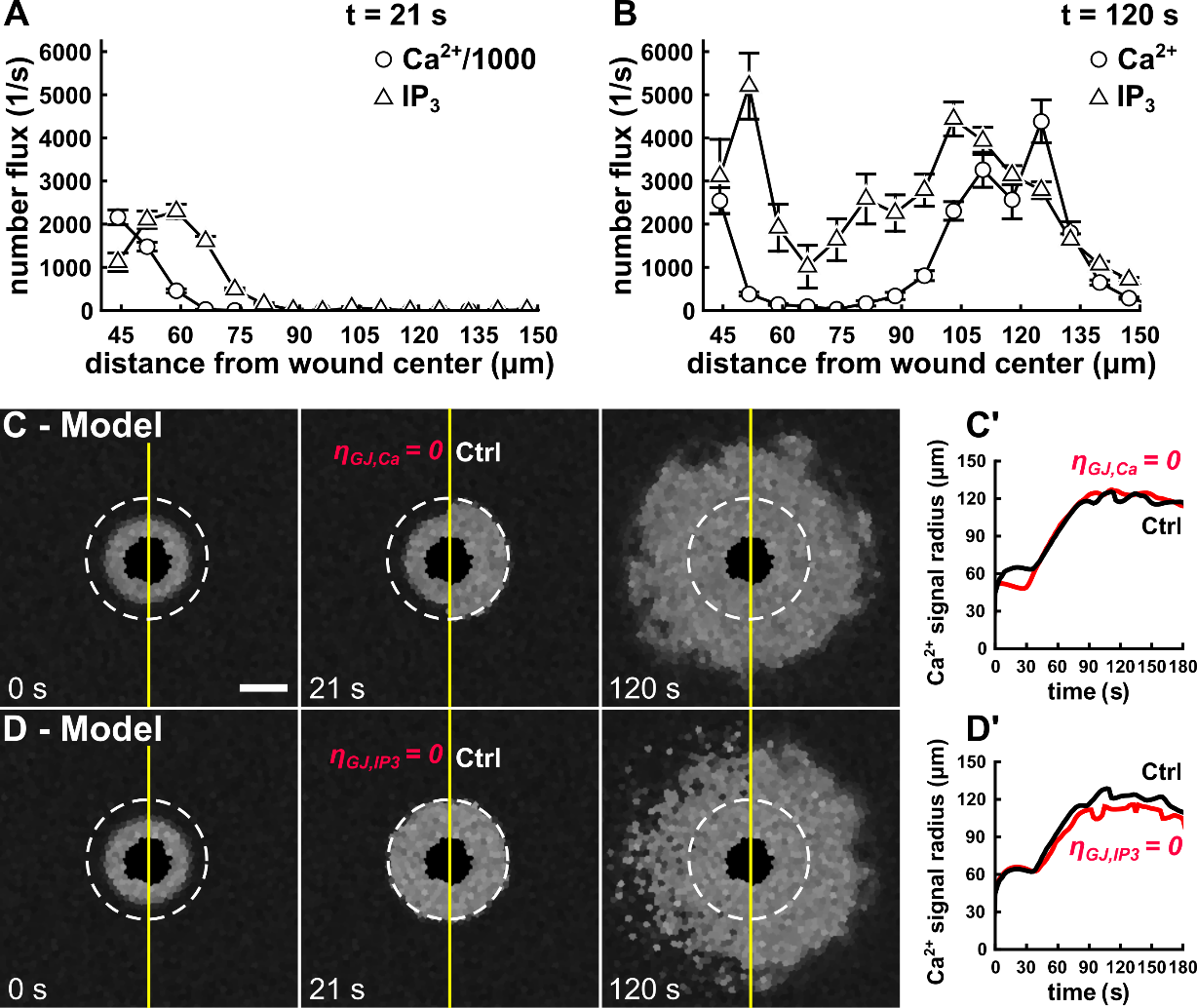

Figure 5: Model demonstrates different roles for gap-junction fluxes of calcium and IP_3_.

(A-B) Quantification of average gap-junction fluxes, binned by distance from the wound center at both the maximum extent of the first expansion (21 s after wounding) and just after completion of the second distal expansion (120 s). Error bars denote standard error of the mean. Note that the calcium fluxes in A have been divided by a factor of 1000 to plot on the same scale as IP_3_ fluxes. (C-D) Model responses after selective *in silico* knockdown of gap-junction fluxes for either calcium (C) or IP­_3_ (D). Scale bar is
50 μm and applies to all images. (C’-D’) Corresponding quantifications of the calcium signal radius as a function of time. See also Movie S13 and Movie S14.
